# The landscape of structural variation in pediatric cancer

**DOI:** 10.1101/2025.04.24.650284

**Authors:** Robert Greenhalgh, Wentao Yang, Samuel W. Brady, Diane A. Flasch, Yanling Liu, Karol A. Szlachta, Liqing Tian, Pandurang Kolekar, Jian Wang, Xin Zhou, Daniela S. Gerhard, Xiaotu Ma, Jinghui Zhang

**Affiliations:** Department of Computational Biology, St. Jude Children’s Research Hospital, Memphis TN, USA; Department of Pharmacy and Pharmaceutical Sciences, St. Jude Children’s Research Hospital, Memphis TN, USA; Office of Cancer Genomics, National Cancer Institute, National Institutes of Health, Bethesda MD, USA

## Abstract

Structural variants (SVs) account for over 60% of the driver variants in pediatric cancer, and in many cases act as the cancer initiating event. To study SVs from a pan-cancer perspective, we analyzed 1,616 pediatric cancer genomes in 16 major cancer types of hematological malignancies (n = 908), brain tumors (n = 183), and solid tumors (n = 525) and compared their profiles to those of 2,203 adult cancers. The SV burden varied ∼100-fold across pediatric cancer types and demonstrated an 8-to 16-fold reduction compared to adult brain and solid tumors but was comparable in pediatric versus adult hematological malignancies. Recurrent SV hotspots occurred uniquely in pediatric acute lymphoblastic leukemias (ALLs) in proximity to RAG-mediated recombination signal sequences (RSS) and disrupted multiple immune-related loci as well as 69 genes, which often involved cryptic RSS sites. By contrast, such hotspots affected only immune-related loci but not driver genes in adult lymphoid cancers. Eight SV signatures extracted from the cohort had varying distributions across cancer types, with clustered translocations reflecting templated insertions in osteosarcoma, and medium-sized deletions (10 kb to 1 Mb) enriched in cancers with RAG-mediated deletions. Intra-patient evolutionary analysis in 13 patients with multiple spatiotemporally distinct samples revealed that RAG-mediated recombination in leukemia and complex rearrangements in solid tumors occurred both early in disease initiation and continuously during later diversification, contributing to clonal heterogeneity. Finally, we found that both driver genes and fragile sites were the two genomic regions most frequently disrupted by SVs. The unique and diverse SV landscapes that emerged from this comprehensive analysis expand the scope of RSS-mediated mutagenesis in pediatric ALL and will be a valuable resource for guiding future functional studies and the design of clinical genomic testing in pediatric cancer.

## Introduction

Genomic structural variants (SVs) occur during the rearrangement of genomic segments, such as through intra-chromosomal large deletions, insertions, or inversions, or inter-chromosomal translocations^1^. Somatically acquired SVs can promote tumorigenesis by: (1) altering the gene dosage in oncogenes via copy gain and in tumor suppressors via copy loss^2^, (2) disrupting critical functional domains in driver genes (e.g. loss of the chromatin-binding domains of *ATRX* in neuroblastoma^3^ and the constitutional dimerization of *FGFR1*^3^ in low-grade gliomas^4^), (3) bringing together two genes to create a fusion oncoprotein^5^, or (4) juxtaposing an active promoter or enhancer to an oncogene^6^. Historically, SVs in the form of chromosomal rearrangements were detected by cytogenetics, and have long been recognized as disease drivers in pediatric leukemia^7^ and solid tumors (e.g. Ewing’s sarcoma^8^ and rhabdomyosarcoma^9^). Their contribution to pediatric cancer development has further expanded over the last decade^10,11^ as whole-genome sequencing (WGS) has enabled the discovery of diverse SV types that disrupt the function or regulation of oncogenes and tumor suppressors.

Mechanistically, SVs are facilitated by double-strand DNA breaks, which may be incorrectly repaired through such processes as non-homologous end joining, and by replication errors in which DNA polymerases aberrantly shift to an alternative template^12^. Double-strand breaks leading to SVs may occur through ionizing radiation^13^, mechanical stress (as in breakage-fusion-bridge cycles^14^), micronucleus formation^15^, enzymatic processes such as RAG1/2-induced recombination at cryptic recombination signal sequence (RSS) sites^16^, or via improper handling of stalled replication forks by DNA replication and repair enzymes^17,18^. The diverse mechanisms underlying SV formation give rise to characteristic patterns or “signatures”, allowing retrospective identification of the processes that may have led to SV formation in individual tumors^19–22^. Though adult somatic SV signatures have been examined in several pan-cancer studies^19,20,22,23^, SV signatures across pediatric cancers, which often have differing genetic drivers from adult cancers^10,11^, are less understood^24–26^.

Here, we assembled a cohort of publicly available paired tumor-normal WGS data from 1,616 pediatric cancer patients to elucidate the landscape of structural variation across 16 major pediatric cancer types. Our analysis reveals diverse mutational processes giving rise to SV alterations in our cohort, identifies genomic features that appear unique to SV mutagenesis in childhood cancer, and provides insights which may inform future functional studies as well as the design of clinical genomic testing in pediatric cancer.

## Results

### Comparison of SV burden in pediatric and adult cancers

To study the SV patterns in pediatric cancer, we aggregated matched tumor-normal WGS from 1,616 pediatric cancer patients (all under 18 years of age) from the St. Jude/Washington University Pediatric Cancer Genome Project (PCGP)^27–33^, the National Cancer Institute’s Therapeutically Applicable Research to Generate Effective Treatments (TARGET) project^11,34–37^, St. Jude’s clinical genomics sequencing programs (including Genomes for Kids, G4K)^38,39^, and our previously studied acute lymphoblastic leukemia samples from Shanghai Children’s Medical Center^40^ (Supplementary Table 1). For landscape-level analysis, a single diagnosis tumor for each patient was analyzed using WGS. Somatic SVs were identified by computational analysis, orthogonal experimental verification, and manual review, which resulted in a final dataset of 40,780 curated somatic SVs (Methods, Supplementary Table 1). Of the 1,616 cancer genomes analyzed, all but 39 (n = 1,577; 97.59%) had at least one curated somatic SV.

The SV burden, defined as the number of somatic SVs per cancer genome, was summarized for the 16 major pediatric cancer types (Figure 1a) in the following three categories for comparison with the corresponding cancer type in adults (Supplementary Table 2): hematological malignancies (n = 908), central nervous system (CNS, hereafter referred as brain tumors; n = 183) and non-CNS solid tumors (hereafter referred as solid tumors; n = 525). The median pediatric SV burden ranged from 1–245, with the ∼100-fold variability among cancer types indicating potential differences in SV mutational processes. Notably, amongst the cancer types with the highest SV burden, osteosarcoma (OS, median 245 SVs), adrenocortical cancer (ACT, 51.5 SVs) and high grade glioma (HGG, 37 SVs), had the highest *TP53* mutation rate in our cohort with prevalences of 91%, 84% and 63%, respectively^32,41,42^. Despite also having a high SV burden (median 34 SVs), rhabdomyosarcoma (RHB) was an exception to this finding, as this solid tumor rarely harbors *TP53* mutations (<12% prevalence). The lowest SV burden was found in acute myeloid leukemia (AML), low grade glioma (LGG), Ewing’s sarcoma (EWS), rhabdoid tumor (RT), and Wilms tumor (WT), each with a median SV burden ≤5. Compared with the adult cancer SV data compiled by the Pan-Cancer Analysis of Whole Genomes (PCAWG) consortium^20^, the median SV burden in pediatric brain (5) and solid tumors (9) showed an 8–16-fold reduction (Figure 1a, Supplementary Figure 1), which was consistent with the reduced somatic single nucleotide variant (SNV) and small insertion/deletion (indel) mutational burden reported in prior pan-cancer analyses^10,11,43^. By contrast, the median SV burden of pediatric hematological malignancies (7) was comparable to that of their adult counterparts (7).

**Figure 1.**
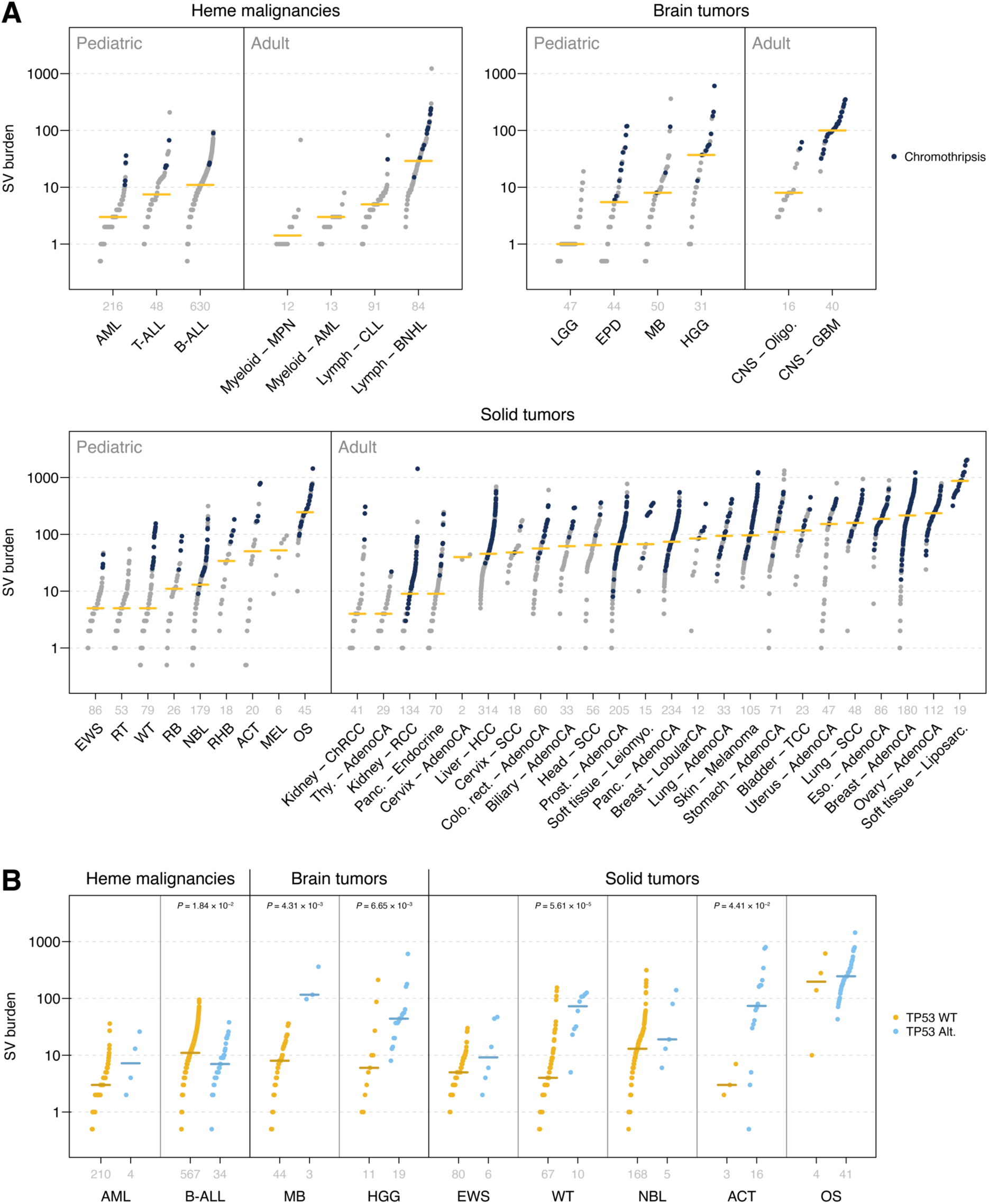
Somatic SV burden in pediatric and adult cancers. (**a**) Number of SVs per tumor in hematological malignancies (top left), brain tumors (top right), and solid tumors (bottom). Pediatric tumor types and adult cancer types from the same category are shown at left and right, respectively, and are labeled with abbreviations used in this study (see Abbreviations) and by PCAWG. Sample numbers per cancer type are shown in gray above each cancer label. Each data point represents one cancer sample, and horizontal yellow lines represent the cancer median. Dark blue dots represent cancer samples with chromothripsis. (**b**) Comparison of SV burden in *TP53*-wildtype versus *TP53*-mutant samples in pediatric cancer. Similar to (a) except that each pediatric cancer type is split into *TP53* wildtype or altered samples. Both germline and somatic *TP53* alterations were considered. Only cancers with at least three *TP53*-altered samples were analyzed. Significant *P*-values by two-sided Wilcoxon rank-sum tests are listed.

Within each pediatric cancer type, SV burdens varied substantially, with the variation exceeding 100-fold in eight of the 16 cancer types (i.e. T-ALL, EPD, MB, HGG, WT, NBL, ACT, OS, see Abbreviations). In line with the SV burden, complex SVs with multiple genomic locations involved in a single rearrangement event (Methods) were found to differ substantially by cancer type (Supplementary Figure 2a). To evaluate one potential source of this high variability, we examined the prevalence of chromothripsis in our data set, which can generate multiple SVs through a single catastrophic event. Chromothripsis was identified in 1.5%, 13.1%, and 14.9% of hematological malignancies, brain tumors, and solid tumors, respectively (Figure 1a, Supplementary Figure 2b). Indeed, there was a 1.1–25.1-fold increase in the median SV burden in chromothripsis-positive samples within each cancer type (Supplementary Figure 2b). Next, we examined the effect of *TP53* mutation on SV burden within each cancer type. In brain and solid tumors, an elevated SV burden was found in *TP53*-mutated samples, consistent with prior studies^10,44^ (Figure 1b). By contrast, the SV burden in hematological malignancies was not significantly increased by mutated *TP53* (Figure 1b). Finally, we analyzed the effect of age at diagnosis on SV burden. Positive correlation was found in six (ACT, B-ALL, NBL, RB, T-ALL, and WT; Supplementary Figure 3) of the 16 pediatric cancer types, in contrast to two (melanoma and prostate) of the 27 adult PCAWG tumor types (Supplementary Figure 4). Notably, the pediatric cancers with the strongest age/SV correlation were ACT (*R* = 0.737) and RB (*R* = 0.728); both cancers are frequently caused by germline mutations in cancer predisposition genes (*TP53* or *RB1*, respectively^42,45^).

### Regions with recurrent SV breakpoints

A striking pattern of recurrent SV breakpoints in multiple cancer genomes emerged when we used `GenomePaint`^46^, a visualization tool for diverse types of DNA variants, to examine SVs that disrupted driver genes. *RB1*, a pan-cancer tumor suppressor disrupted by intra-or inter-chromosomal SVs in pediatric brain tumors (HGG), solid tumors (RB and OS), as well as leukemias (B-ALL, T-ALL and AML), is one such example (Figure 2a). Specifically, an SV breakpoint hotspot in intron 17 of *RB1* is connected to either a breakpoint hotspot within the neighboring *RCBTB2* gene or one in the intergenic region upstream of the long-noncoding RNA *LINC00462*, resulting in two recurrent deletions that truncate the C-terminus of *RB1* (Figure 2a). These three SV hotspots were exclusive to B-ALL and T-ALL, as no such hotspot was found in other pediatric cancer types (including RB), nor in adult cancers. They involve 16 ALL cases, most of which harbor kinase fusions such as *BCR-ABL1* or *BCR-ABL1*-like (e.g. *EPOR*, *CRLF2*). To investigate if these three hotspot were caused by RAG-mediated recombination — a mechanism previously reported to generate deletions in the *ETV6-RUNX1* subtype as well as create deletion hotspots at *CDKN2A* loci in several lymphoid leukemia cell lines^47,48^ — we examined the sequence context flanking the recurrent breakpoint hotspots, which revealed the presence of the RAG heptamer (CACAGTG) located within 20 bp of all three SV hotspot breakpoints (Figure 2b).

**Figure 2.**
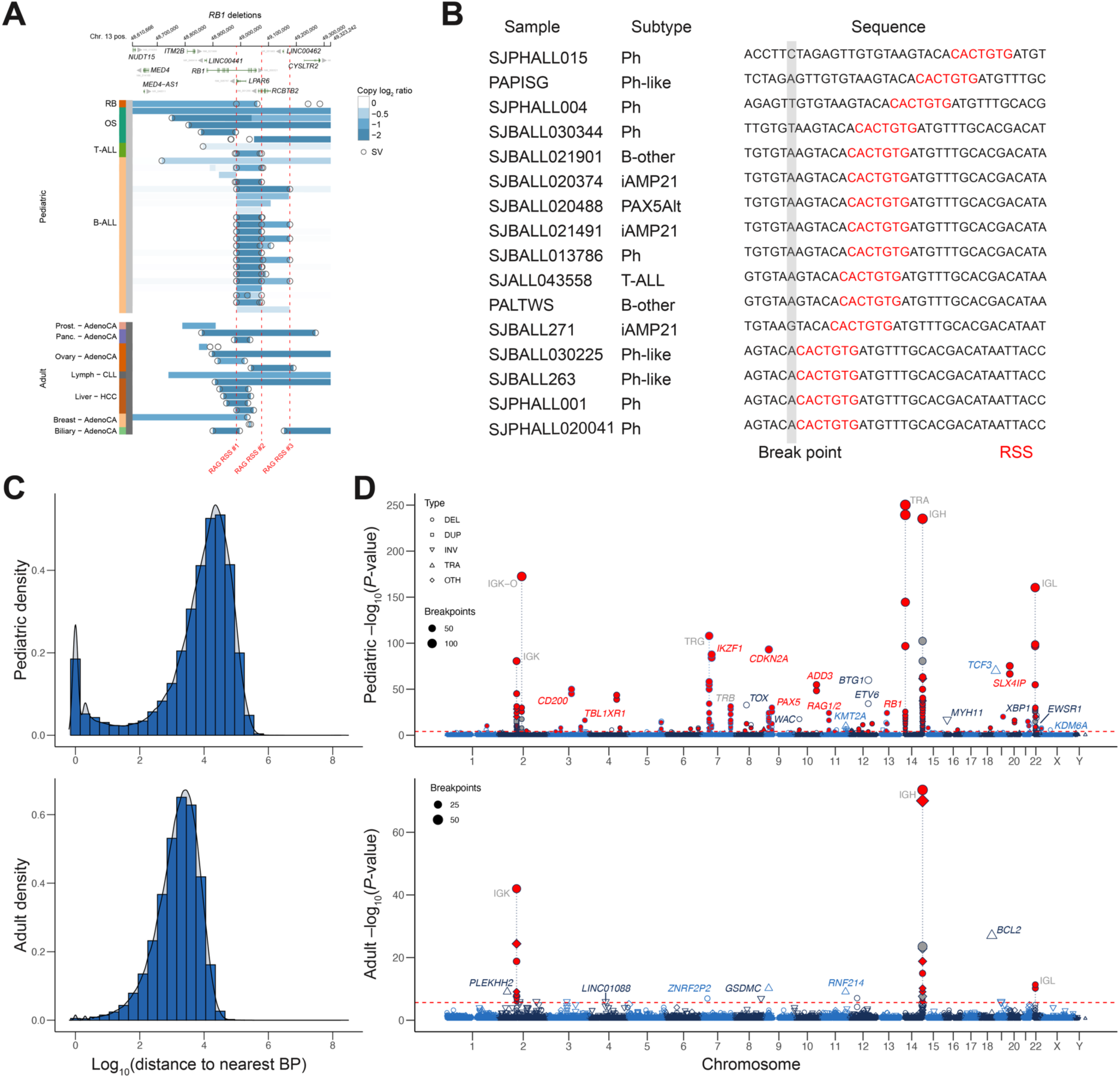
Genomic locations of recurrent SV hotspots. An illustration of SV hotspots at *RB1* locus in pediatric ALL is shown in (**a**) and (**b**). (**a**) Copy number and SV profiles for pediatric (top) and adult (bottom) cancers with *RB1* deletions. Each row represents one sample, with cancer type indicated on the left. Circles denote the locations of SV breakpoints while blue coloring indicates copy loss. Vertical red dotted lines indicate the locations of the three hotspots in ALL, all of which are associated with RSS sites. (**b**) RSS heptamer sequences in close proximity to the hotspot labeled as “RAG RSS1” in (a) present in representative ALL samples with diverse subtypes. (**c**) Density plots showing the distribution of the nearest breakpoint distance across pediatric (top) and adult (bottom) cancers. (**d**) Manhattan plots of genome-wide SV hotspots in pediatric (top) and adult (bottom) cancers. Chromosomes are shown in alternating blue and black colors. An FDR of 0.05 is indicated by the red dashed horizontal line. Each point represents one SV hotspot, with the shape and size indicating the dominant (≥75%) SV type (labeled with abbreviations: DEL (deletion), DUP (duplication), INV (inversion), TRA (translocation), OTH (other, i.e. no dominant SV type)) and number of samples in the hotspot, respectively. Red coloring indicates hotspots where at least 75% of breakpoints are within 20 bp of an RSS site. Immune related hotspots are labeled in gray and denoted with a dotted gray line.

To analyze this pattern at genome-scale, we first plotted the inter-tumor SV distance distribution in our pediatric cancer cohort and compared it to that of the PCAWG adult cancer data (Figure 2c). While a single peak at ∼2,000 bp was observed in adult cancer, a bi-modal distribution was found in pediatric cancer with dual peaks at <100 bp and ∼12,000 bp. The peak at ∼12,000 bp, reflecting the reduced SV burden in pediatric cancer, corresponds to the primary peak at 2,000 bp in adult cancer. By contrast, the peak at <100 bp is unique to pediatric cancer and not caused by chromothripsis, as it is absent when analyzing chromothripsis-positive pediatric cancer genomes (Supplementary Figure 5a). We further examined the SV distribution within each cancer type and found that the bimodal distribution was most prominent in lymphoid leukemias which include B-ALL and T-ALL (Supplementary Figure 5a).

We next examined genomic regions that exhibit significant enrichment of SV occurrence and identified 272 hotspots of recurrent SV breakpoints that gave rise to the unique peak in pediatric cancer (Figure 2d). The vast majority of these SV hotspots (85.3%; 232 of 272) result in deletions. The SVs within the hotspots were contributed by B-ALL (88.6%), T-ALL (3.2%), AML (4.0%), and EWS (3.3%). Hotspots contributed by AML and EWS were driven by gene fusions in the corresponding introns of *KMT2A, MYH11* and *EWSR1*. Altogether, only 9 hotspots (3.3% of the total) were contributed by inter-chromosomal translocations resulting in the gene fusions of *TCF3*, *KMT2A, MYH11*, and *EWSR1* (Supplementary Figure 6).

Motif analysis by `HOMER`^49^ identified RSS as the sole significant motif across all hotspot regions, indicating RAG-mediated recombination contributed to the formation of these hotspots (Supplementary Figure 5b, left). Almost 60% of the hotspots (161 of 272) were in regions involved in immune gene rearrangement; 88.1% (141) of which were associated with RAG-mediated RSS sites, consistent with prior knowledge of the role of RAG in catalyzing the rearrangement of the variable (V), diversity (D), and joining (J) DNA elements of antigen receptor genes^50^.

Amongst the remaining hotspots unrelated to immune gene rearrangements, the majority (77.5%; 86 of 111) were found within intragenic regions (69 genes in all), similar to the *RB1* example in Figure 2a. Predicted RSS sites were associated with 59.5% (66 out of 111) of these non-immune hotspots, and significant enrichment of RSS heptamer sequence was evident (*P* = 10^-20^; Supplementary Figure 5b, right). We were able to replicate SV hotspots previously reported in known leukemia driver genes such as *CDKN2A*, *BTG1*, *PAX5* and *TCF3-PBX1* (Figure 2d, Supplementary Figure 6). Importantly, many of these SV hotspots have not been previously reported, including SV hotspots involving well-established driver genes such as *RB1*, *CRLF2*, *KDM6A*, and *LEF1*. Even for loci extensively studied for RAG-mediated recombination such as *CDKN2A*, new hotspots were identified in our cohort (Supplementary Figure 7, red lines). The majority of B-ALL (85.9%) and T-ALL (68.8%) contain at least one SV within the hotspot regions and the vast majority (88.6%) of SVs within the hotspots were from B-ALLs due to the large sample size and high prevalence of this subtype. *RAG1* and *RAG2* expression levels in B-ALLs that harbored non-immune hotspots were elevated compared to that of the negative cases (Supplementary Figure 5c), indicating the effect of RAG activity on generating recurrent SVs in these regions.

Performing the same analysis on 275,430 SVs from adult PCAWG samples yielded 39 recurrent SV hotspots, demonstrating that such events were rare in the PCAWG adult cohort despite a 6.75-fold increase in the total number of SVs (Figure 2d, bottom). The majority (61.5%) of PCAWG SV hotspots were in immune regions and found in lymphoma samples, and *BCL2* is the only significant genic region which is involved in *BCL-MYC* translocation in lymphoma (Supplementary Figure 8).

### Differential selection for RSS sites at immune and non-immune regions

The SV hotspots at non-immune regions raised an interesting question on whether they were selected to function as potential recombination sites, which can be evaluated by the Recombination Information Content (RIC) score, a measure of the potential functionality of an RSS site based on mutually correlated positions^51^. As expected, at immune loci like TRA and IGK, sites exhibited high predicted recombination efficiency (equating to higher RIC scores; Supplementary Figure 9); in both pediatric (immune median: 18.9; Supplementary Figure 10) and adult (adult immune median 21.0; Supplementary Figure 11) hematological malignancies. In non-immune regions, however, this pattern was absent (pediatric non-immune median: 3.74; Supplementary Figure S9). While cryptic RSS sites in or near genes like *CDKN2A* and *RB1* were not expected to have RIC scores equivalent to immune loci, the sites linked to pediatric RAG-mediated deletion events were not even found to be the best predicted RSS sites in their respective regions — indeed, the most prominent site in *RB1*, host to breakpoints from 23 samples, barely passed the threshold for identification as a cryptic RSS site (RSS score: 0.20; Supplementary Figure S10d, red arrowhead).

### SV signatures in pediatric cancer

To gain further insight into the SV mutagenesis processes in pediatric cancer, we classified SVs into one of 32 SV types^19,21^ (Figure 3a). First, intra-chromosomal SVs were classified as deletions, duplications, or inversions based on the orientation of the two breakpoints, while inter-chromosomal translocation SVs were classified as translocations. Intra-chromosomal SVs were further subclassified by the distance between the two breakpoints into five bins: 1–10 kb, 10–100 kb, 100 kb–1 Mb, 1–10 Mb, or >10 Mb. Finally, SVs were classified by whether they clustered near other SV breakpoints within the same tumor sample; note this was analyzed for each individual tumor sample as an intra-tumor feature. Following SV classification, we performed *de novo* SV signature extraction using `SigProfilerExtractor`^52,53^, a tool which utilizes a non-negative matrix factorization (NMF) approach to identify mutational signatures.

**Figure 3.**
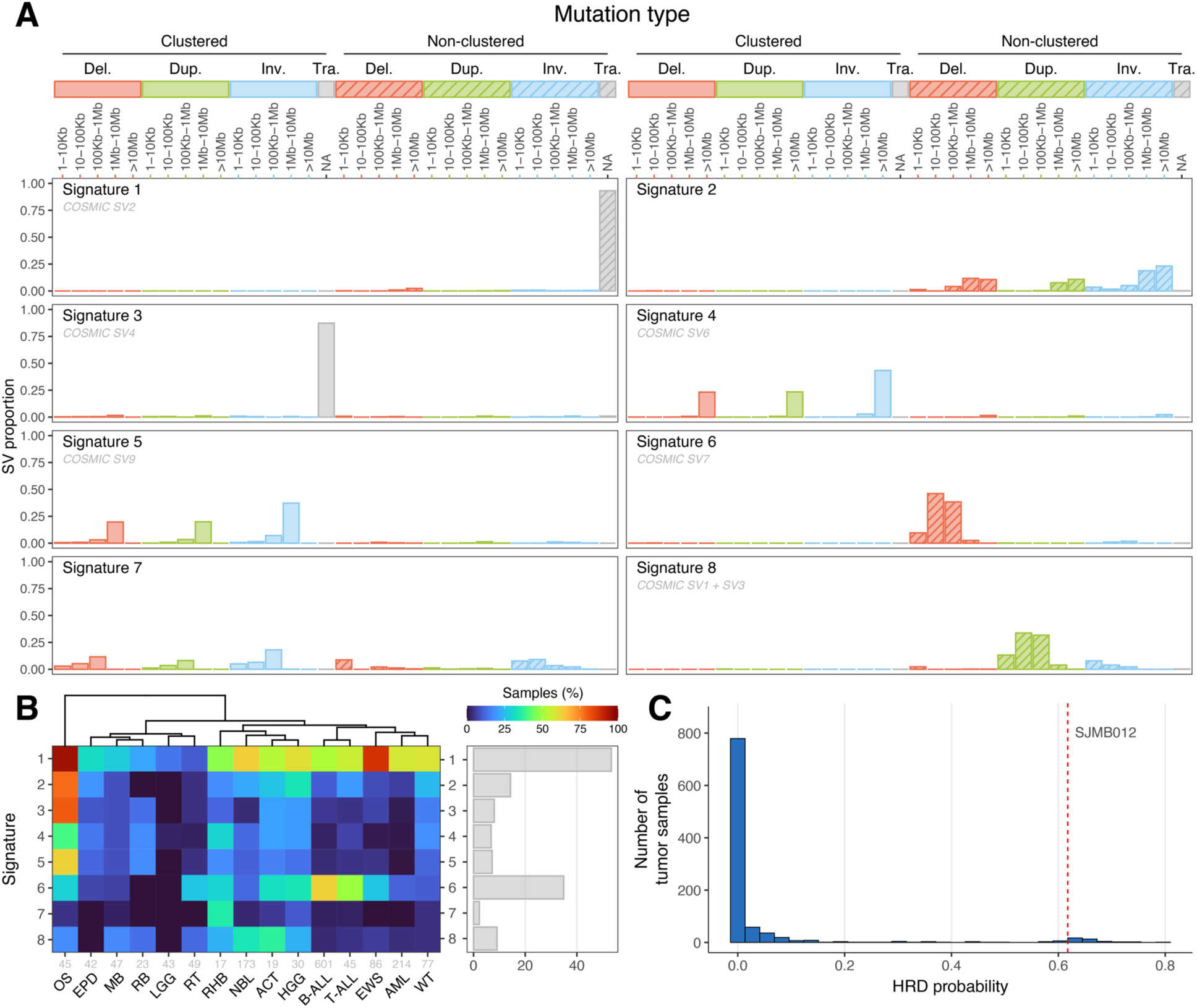
Somatic SV signatures in pediatric cancer. (**a**) Bar plots showing the distributions of the 32 SV types in all eight SV signatures extracted from the pediatric cancer cohort, with non-clustered signatures represented by patterned bars. Signatures with matches to the COSMIC database have their corresponding COSMIC signature(s) listed in gray below the signature name. (**b**) Heatmap showing the percentage of cancer samples with each SV signature shown in (a) for all cancer types with at least 15 samples in the dataset. The total number of cancer samples for each cancer type is shown in gray. The bar plot at right shows the percentage of all pediatric cancer samples exhibiting each signature. (**c**) Distribution of HRD probability in pediatric cancer. The red line denotes SJMB012, an HR-deficient medulloblastoma with bi-allelic loss of *BRCA2* caused by compound heterozygosity mutations in the germline^57^.

A total of eight SV signatures (Figure 3a) were identified, each with varying prevalence across cancer types (Figure 3b). For each cancer type, the association between a given SV signature and chromothripsis was also evaluated, showing a general enrichment of signatures representing clustered SVs in chromothripsis. (Supplementary Figure 12). Six of the eight signatures extracted were similar (cosine score ≥0.80) to one or more of the ten previously reported SV signatures found in the COSMIC database^22^, while two (signatures 2 and 7) had cosine scores below this threshold (Figure 3a).

Signature 1, which matched COSMIC signature SV2 and was defined by non-clustered translocations, was present in most pediatric cancers (53.3%) and enriched in OS (detected in 95.6% of samples) and EWS (87.2%). Signature 2, which had no match in the COSMIC database, was comprised of medium to large non-clustered deletions, duplications, and inversions, and was most prevalent in OS and to a lesser degree HGG.

Signature 3, which represents clustered translocations and corresponds to COSMIC signature SV4, was most prevalent in OS but intriguingly, not associated with the chromothripsis status of OS. A deep look into one of the samples with the highest inter-chromosomal SV burden (SJOS013 with >750 translocation SVs) revealed a pattern resembling a “cycle or chain of templated inter-chromosomal insertions”, originally described by the PCAWG group in adult cancers^20^ instead of bona-fide translocation events. An example of inter-chromosomal insertion in SJOS013 is shown in Supplementary Figure 13a, where clusters of 2–4 rearrangements within 1–2 kb were detected in five regions across three chromosomes.

Signatures 4 and 5, defined by large (>10 Mb) and medium-large (1 Mb to 10 Mb) clustered intra-chromosomal SVs, respectively, were most prevalent in OS. Both signatures had matches in the COSMIC database (SV6 and SV9, respectively), and were also associated with chromothripsis status in multiple cancer types (Figure S12), with the most significant association found between signature 4 and chromothripsis in NBL.

The remaining signatures did not exhibit significant association with chromothripsis status. Signature 6, corresponding to COSMIC SV7, was dominated by medium-sized deletions (10 kb to 1 Mb) and was most abundant in B-ALL (62.6%) and T-ALL (48.9%), suggesting that RAG1/2-mediated deletions may contribute to this signature^47^. Indeed, the deletion size distribution of recurrent SV hotspots in regions with the RSS motif matches this projected range (Supplementary Figure 13b). Signature 7 consisted of a mixture of small-medium sized clustered and non-clustered intra-chromosomal events and was enriched only in RHB; due to the low prevalence of this signature in our dataset (2.3%), as well as its failure to match any known COSMIC signature (cosine similarity of closest match: 0.531), we suspect this signature may be an artifact of the `SigProfiler` analysis. Signature 8, corresponding to a combination of COSMIC SV1 and SV3, involved non-clustered duplications of intermediate size (10 kb to 1 Mb) and was most common in ACT (36.8%) and NBL (33.5%).

As SV signatures are known to be associated with the presence of *BRCA1* or *BRCA2* mutations or promoter hypermethylation status^19^, this prompted us to perform an analysis on homologous recombination deficiency (HRD). This process, which is one of the proposed mechanisms underlying COSMIC SV3 (and therefore our signature 8), is mediated by the *BRCA1/2*, *RAD51* and *PALB2* genes and confers selective sensitivity to compounds such as PARP inhibitors^54^. We ran `CHORD`^55^, a random forest classifier trained on pan-cancer WGS data, to predict HRD based on somatic SNVs, indels, and SVs. HRD was predicted in 2.6% of the cases (Figure 3c, Supplementary Table 3), which included SJMB012, a MB with bi-allelic loss of *BRCA2* caused by compound heterozygosity mutations in the germline^56,57^.

Signature analysis of the adult cohort yielded a total of ten signatures, eight with matches to the COSMIC database (Supplementary Figure 14a). All COSMIC signatures identified in the pediatric cohort were found in the adult analysis; COSMIC SV8, which was absent from the pediatric analysis, was also identified. The two signatures without COSMIC matches, adult signatures 2 and 6, consisted of multiple non-clustered events of varying sizes and very small non-clustered deletions, respectively. Notably, while adult signature 4, which corresponded to COSMIC SV7/pediatric signature 6, was also found to be enriched in adult hematological malignancies, it was found to have even stronger enrichment in esophageal and colorectal adenocarcinomas, which do not have RSS-associated hotspots, indicating potential alternative mechanisms for COSMIC SV7 (Supplementary Figure 14b).

### Temporal evolution of SVs

To explore whether SV formation is involved in tumor clonal evolution, we analyzed 13 pediatric cancer patients with multiple spatiotemporally distinct WGS samples (and patient-derived xenografts in some cases). These included three OS patients with 2–6 samples each with matching patient-derived xenograft (PDX) mouse models analyzed for two patients (Figure 4a, b, Supplementary Figure 15), two RHB patients with 2 and 3 samples each (Supplementary Figure 16), two NBL patients with matched diagnosis and relapse samples (Supplementary Figure 17), and six B-ALLs with matched diagnosis and relapse samples (Figure 4c). To understand the evolutionary pattern of SVs in each patient, we grouped SVs based on their presence or absence in each tumor sample. SVs detected in all samples of a patient were considered “truncal” (early events), those detected in multiple, but not all samples were considered “shared” (intermediate events), and those present only in a single sample were considered “private” (late events).

**Figure 4.**
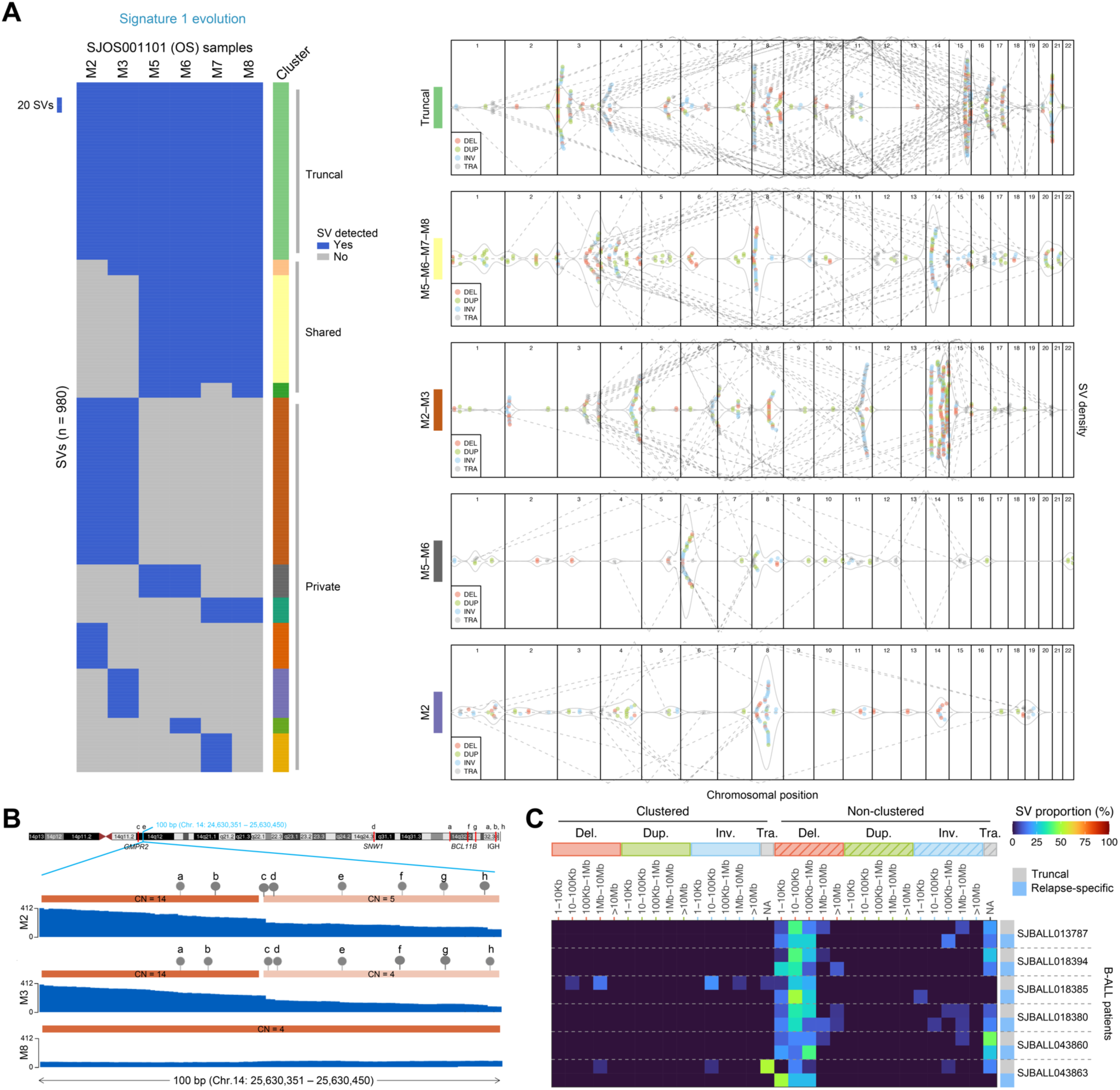
SV evolution in multi-sample osteosarcoma and B-ALL patients. (**a**) Left, heatmap showing presence (blue) or absence (gray) of SVs in six metastatic osteosarcoma samples obtained at autopsy from one patient. Each column represents one sample, and each row a single SV. SVs are grouped by presence or absence in each sample into distinct evolutionary categories, as shown by the colors at right. Right, beeswarm plots showing the SV density (gray line) for selected SV evolutionary groups (matched by color to the groups shown in heatmap at left) across the genome. Each point represents one SV breakpoint with the SV type color-coded using the same scheme as Figure 3. Dotted gray lines show the joining of translocations. (**b**) Detailed view of an SV cluster in the M2-M3 group in (a) which has eight SVs within 100 bp connected to seven distinct genomic regions of chromosome 14. (**c**) Comparison of SV signatures at six B-ALL samples at diagnosis and relapse. Heatmap showing the proportion of SVs falling into each of the 32 SV types. Each row represents the truncal (present at both diagnosis and relapse) or relapse-specific SV profile for one patient.

All three OS patients analyzed harbored somatic *TP53* inactivation and had numerous translocations occurring continuously through evolution, including in PDX models, which arose from chromothripsis or templated insertions (Supplementary Figure 18). This pattern was visualized by plotting SV density across the genome and connecting translocation breakpoints with dotted lines as shown in Figure 4a and Supplementary Figure 15 (right panels). For example, in SJOS001101 (a patient with six high-purity samples acquired at autopsy from multiple metastases in the left and right lungs), truncal translocations primarily affected chromosomes 4, 11, 16, and 17, while later shared translocations (in samples M2 and M3; group M2-M3) occurred on chromosomes 3, 6, and 11. Shared SVs, which primarily occurred in metastatic samples collected at left (i.e. M2-M3) or the right (i.e. M5-M6-M7-M8) lung section, matched the shared SNV/CNV profile reported previously for this patient^58^. The high SV density on chromosome 14 unique to the shared SVs in M2-M3 occurred in short segments (most <1 Kb) with high-level amplification (>10 copies), and the breakpoints were connected to multiple dispersed genomic regions. An example of eight SVs within 100 bp connected to seven distinct regions on chromosome 14 is shown in Figure 4b and Supplementary Figure 18. Interestingly, the read count for these SV breakpoints in the M2 and M3 samples were highly concordant (Supplementary Figure 18c), indicating a possibility that these SVs and amplicons arise from a single, late event.

A similar pattern of continued SV evolution can be found in the two embryonal RHB patients with matched diagnosis and relapse samples (Supplementary Figure 16). For example, for case SJRHB012, clustered SVs were found on chromosome 12 in three distinct regions: p13.1–13.2, q15 and q24. Truncal SVs at q15 caused the formation of a double-minute amplicon with ∼100 copies targeting the *MDM2* oncogene in tumor samples acquired at diagnosis and relapse. By contrast, the SV clusters at 12p13.1–13.2 were distinct in diagnostic and relapsed samples, which joined genomic fragments at 12q15 and 22q11 (Supplementary Figure 16b–e), respectively. These resulted in distinct *MDM2* amplicons at diagnosis and relapse exhibiting differential gene expression of affected driver genes (i.e. *MDM2* and *ETV6* in Supplementary Figure 16c, d).

SV temporal evolution was also observed in NBL, a solid tumor with a lower SV burden compared to OS and RHB. Of the two NBL patients with matched diagnosis and relapse samples (Supplementary Figure 17a, b), complex SVs involving multiple genomic loci were rare except for a diagnostic-specific SV cluster connecting 6q and 7p in one of the cases (i.e., SJNBL188, D-private, Supplementary Figure 17b). An integrated view of SV and CNV data revealed a chromothripsis event involving ∼30 SVs accompanied by deletions (manifested as two-copy regions on 7p due to a one-copy gain) ranging from <100 bp to 19 Mb (Supplementary Figure 17c). *ARID1B*, a known neuroblastoma driver gene^59^, was the prime target as its first four exons were truncated by this event (Supplementary Figure 17d, left). Interestingly, two common fragile site (CFS) genes, *MAD1L1* and *SDK1*^60^, were both disrupted by this event (Supplementary Figure 17d, right), indicating chromosome breakage at CFS genes may contribute to this complex rearrangement.

In the six B-ALL patients with matched diagnosis and relapse samples (and at least 20 truncal SVs and 20 relapse-specific SVs), non-clustered deletions under 1 Mb in size (i.e. the RAG1/2-associated pediatric signature 6) occurred both in truncal and relapse-specific SVs in similar proportions. This suggests that RAG1/2-induced deletion may be a constant feature of B-ALL evolution and thus may fuel leukemia heterogeneity and relapse (Figure 4c).

### Genes disrupted by SVs

When considering SVs located in gene coding regions, a total of 10,262 genes were disrupted by at least one somatic SV in pediatric cancer. Considering the most frequently SV-disrupted genes with >1% prevalence in our cohort, we found 100 genes that could be classified into the following groups (Figure 5a, Supplementary Figure 19): (1) known driver gene (n = 43); (2) passenger or fusion partner of an SV affecting a driver gene (n = 17); (3) rearrangement of T-cell receptor or immunoglobulin gene (n = 8); (4) fragile site (n = 22); (5) members of the neuroblastoma breakpoint family (*NBPF*, n = 4); and (6) gene of unknown status (n = 6). The most frequently affected gene, *LOC105370401* (Supplementary Figure 19), is a non-coding gene located in the T-cell receptor locus on chromosome 14, and thus likely reflects normal recombination in ALL samples^23^. The 22 fragile-site genes were comprised of large genes (>500Kb) with SV breakpoints in multiple cancer subtypes; many belong to well-characterized common fragile site (CFS) genes such as *CNTNAP2*^61^*, DLG2*^62^, *DMD*^62^, *PTPRD*^63^, *AUTS2*^64^, and *MACROD2*^65^. Among all tumor subtypes, osteosarcoma had the highest frequency of SVs in CFS genes (Figure 5b), consistent with its overall high SV burden in pediatric cancer.

**Figure 5.**
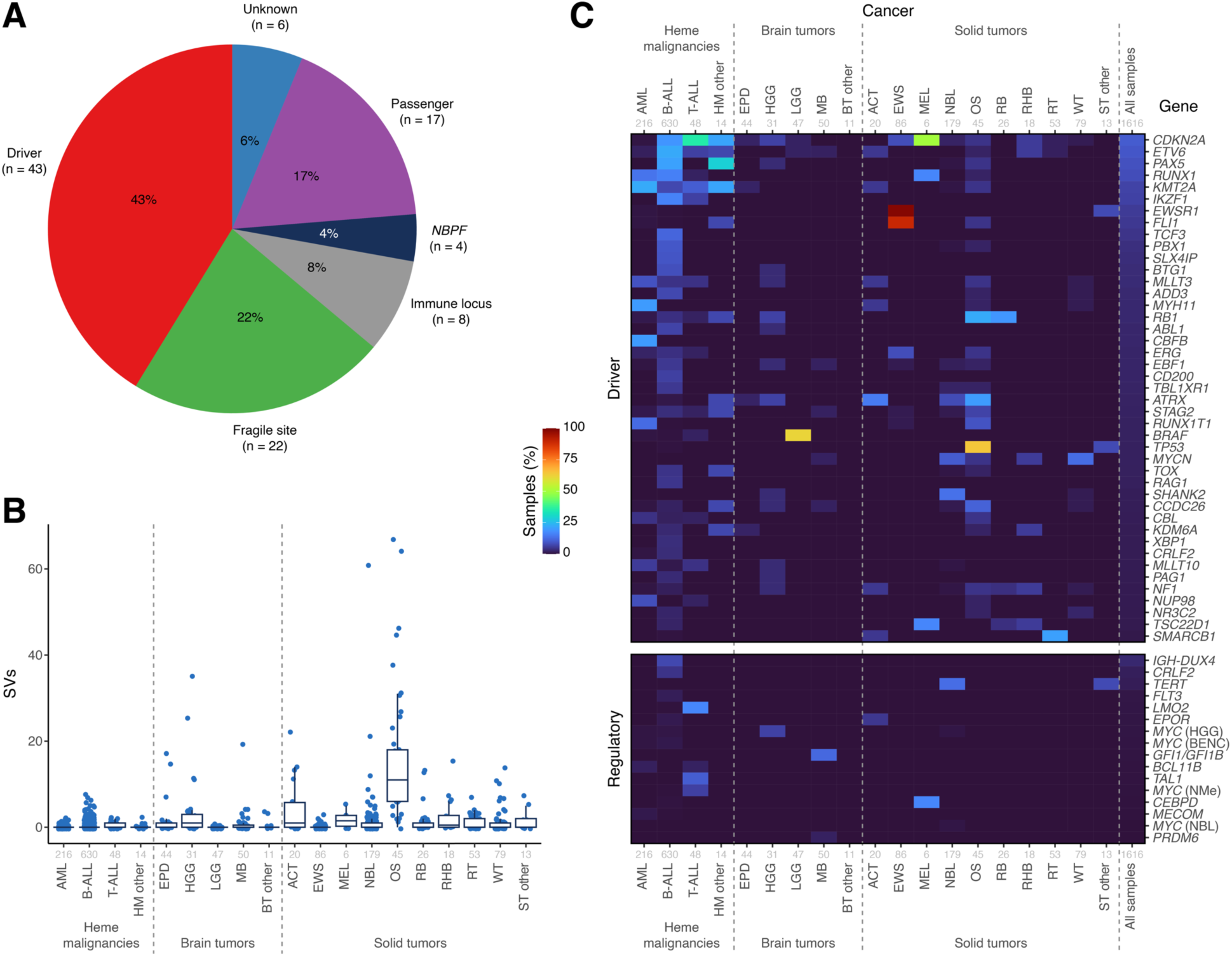
Frequency of genic and regulatory SVs. (**a**) Pie chart showing the types of gene coding regions impacted by SVs in at least 1% of the pediatric cohort. (**b**) Number of somatic SVs in common fragile site genes by cancer type. (**c**) Heatmap showing the percentage of samples in each pediatric cancer type (column) with somatic SVs impacting a driver gene in (a) (row) or disrupting a regulatory non-coding region. A gene was considered impacted in a sample if an SV breakpoint fell within the gene coding region (top) or the regulatory non-coding regions (bottom) with the heatmap showing their prevalence in the relevant cancer types. The number of samples in each cancer type is shown in gray text.

Driver genes mostly frequently disrupted by SVs (Figure 5c, top) were comprised primarily of drivers specific to certain cancer types; this includes *ETV6*, *PAX5*, *RUNX1*, and *KMT2A*^66^ in leukemia, *EWSR1-FLI1* fusion in Ewing’s sarcoma, *TP53* in osteosarcoma, and *BRAF* fusion in low grade gliomas. Pan-cancer SV target genes include genes involved in the cell cycle (*CDKN2A*, *RB1*), cohesion complex (*STAG2*), telomere maintenance (*ATRX*), RAS signaling (*NF1*), and a non-coding RNA gene involved in cMYC regulation (*CCDC26*).

We also analyzed intergenic regions to ascertain the frequency of SVs affecting enhancers or promoters known to cause aberrant overexpression of oncogenes (Figure 5c, bottom). This can be achieved by juxtaposing an active enhancer or promoter to an oncogene (e.g. *IGH-DUX4* rearrangements^28^) through translocation, deletion, or inversion, or by increased enhancer activity through copy gain (e.g. the *NOTCH1 MYC* enhancer (NMe) or blood enhancer cluster (BENC) enhancer regions near the *MYC* gene^46^). The genes most frequently affected by intergenic SVs were *IGH-DUX4* (7.3% of B-ALL)^28^, *CRLF2* (3.5% of B-ALL)^67^, *TERT* (12.8% of NBL)^68^, *FLT3* (1.6% of B-ALL)^69^, and *LMO2* (16.7% of T-ALL)^70^. *MYC* SVs affecting MYC regulator regions including BENC and NMe were found in 15 cases (0.9% of the cohort) of AML, B-ALL, HGG, NBL, and T-ALL.

## Discussion

In this first comprehensive analysis of structural variation in pediatric cancer, we characterized the SV landscape in 1,616 cancer genomes, unveiling a ∼100-fold variability in SV burden across 16 major cancer types. The SV burden of OS, ranked the highest in pediatric cancer, was equivalent to that of ovarian cancer, the second highest tumor type in adult cancer (Figure 1a). By contrast, when considering the mutation rate based on single nucleotide variants (SNVs), OS was 5–10 folder lower than ovarian cancer. This suggests that the mutagenesis processes giving rise to SVs are distinct from those of point mutations. The general perception that pediatric cancer has a lower mutational burden than adult cancer was reinforced by the SV burden in pediatric brain and solid tumors, which were indeed 8–16-fold lower than their adult counterparts. However, contrary to this expectation, the SV burden of hematological malignancy (HM) in pediatric cancers was equivalent to that of adults (Supplementary Figure 1), indicating a potential SV mutagenesis mechanism unique to pediatric HM. Within each tumor type, SV burden also exhibited high variability, and an elevated burden was associated with *TP53* mutation status for brain and solid tumors, though not for HM (Figure 1b). A possible explanation for this is that acute leukemias have a relatively small cell size and high nuclear-to-cytoplasm ratio^71,72^, which might spatially constrain DNA copy gains and hence limit certain SVs. Chromothripsis, detected in 0–53.3% of the genomes of these 16 tumor types, further contributed to a higher SV burden in each tumor type (Supplementary Figure 2b).

We identified 272 regions with recurrent SV hotspots present in our pediatric cohort, while such a pattern rarely occurred in adult cancer (Figure 2). Motif analysis coupled with their predominant presence in ALL indicated RAG-mediated recombination as the most plausible mechanism, consistent with previous studies that focused on a small number of genomic loci or ALL subtypes such as those with *ETV6-RUNX1* fusions^47^. In this study we found that the majority of B-ALL (85.9%) and T-ALL (68.8%) contained at least one SV within these regions, and that most of these hotspot regions were not reported previously (Figure 2a). Thus, RAG-mediated recombination impacted nearly all subtypes of pediatric B-lineage and T-lineage ALL (Figure 2a), contributing to a broader spectrum of drivers than previously recognized; it may also play a role in elevating the pediatric SV burden to a level equivalent to that of adults. Interestingly, RSS sites at non-immune hotspot regions are often cryptic with significantly lower potential for RAG-mediated recombination compared to those in the immune regions, suggesting their preference is therefore likely driven by a factor other than sequence conservation, such as chromatin accessibility^73^. The high recurrence of SV breakpoints in major driver genes in pediatric ALL can be leveraged for optimizing the design of clinical genomic assays, as probes targeting these regions can improve the sensitivity and specificity for detecting SVs, which has historically been a challenge for gene panel testing.

Amongst the eight SV signatures identified in our pediatric cohort, non-clustered translocations (signature 1) were the most prevalent, as this is the predominant mechanism to generate gene fusions or truncations, which are the initiating event in many pediatric cancer types. Medium-sized deletions ranging from 10– 1,000 kb (signature 6, corresponding to COSMIC SV7) ranked second with a high prevalence in B-ALL and T-ALL, suggesting a potential connection to RAG-mediated recombination; this range was also consistent with the size distribution of deletions at recurrent SV breakpoints caused by RAG-mediated recombination (Supplementary Figure 13b). To our knowledge, this is the first aetiology proposed for the COSMIC SV7 signature. The enrichment of this signature in adult lymphoid cancer provides further support for our proposed aetiology, although the presence of COSMIC SV7 in several adult solid tumors indicates the possibility of multiple contributing mechanisms.

Our exploratory analysis on SV evolution in 13 pediatric cancer patients with multiple spatiotemporally distinct WGS samples showed that SV evolution followed the trajectory of clonal evolution as defined by point mutations (SNVs and indels). Importantly, complex rearrangements were an integral part of this process, suggesting that they were subjected to the same level of selective pressure during treatment as point mutations. This indicates that at least some of the mutagenesis processes giving rise to SVs persisted throughout tumor evolution (Supplementary Figures 15–17). In this analysis, we also identified a new type of complex SV in OS which can generate amplicons involving many small rearranged genomic fragments (e.g. eight distinct SVs within 100 bp as shown in Supplementary Figure 18). Current WGS data, which is based on short-read sequencing, is limited in depicting the full assembly of such events, which will require future investigation involving the use of long-read sequencing technology to explore the underlying mutagenesis mechanism.

Finally, when examining the regions of the genome most frequently disrupted by intragenic SVs, we found that driver genes were the likeliest to be impacted, representing 43% of the genes disrupted in at least 1% of our cohort (Figure 5). Fragile site genes were the second most common (22%), followed by passenger genes (17%), immune loci (8%), and *NBPF* genes (4%). Though this demonstrates that the targeting of driver genes is indeed the most common source of recurrent SV events across pediatric cancer types, it also highlights the extremely high rate of other genomic alterations caused by these same mutational processes.

Our comprehensive analysis on SV burden, recurrence, signature and driver genes in pediatric hematological malignancies, brain tumors and solid tumors has provided valuable insights into the SV landscape of pediatric cancer. Our exploratory analysis using multiple spatiotemporally distinct WGS samples further demonstrates that SV-based mutagenesis is an ongoing process contributing not only to tumor initiation but also to clonal evolution driven by the selective pressure of therapeutic exposure.

## Methods

### Patient samples and data

Pediatric cancer patient samples were aggregated from existing WGS data from the PCGP^27^, St. Jude Children’s Research Hospital clinical genomics efforts (including Genomes for Kids, G4K)^38^, TARGET^11^, and previously published WGS from Shanghai Children’s Medical Center and other hospitals in China as part of our prior study^40^. Samples were analyzed under informed consent from parents or guardians and under approval from the institutional review board (IRB) of the applicable institution (including the IRBs of St. Jude, COG member institutions, and Shanghai Children’s Medical Center).

Adult cancer SV data was obtained from PCAWG through the ICGC data portal using existing somatic SV calls^20^. Duplicated diagnostic samples from the same patient were removed by keeping the one with the highest number of SVs. Adult SVs were annotated with the `FusionBuilder`^33^ pipeline, and SVs identified as potential duplicates were filtered out.

### SV identification in pediatric cancer samples

984 samples were profiled by Illumina WGS, and `CREST`^74^ v.1.0 was used to identify somatic SVs following alignment to the hg19 reference with `BWA`^75^ v.0.5.9. SVs generated from 256 PCGP samples and all SCMC samples (n = 98) were experimentally validated by targeted capture sequencing which involved designing probes targeting the SV junction followed by NGS sequencing in both tumor and normal samples. Unvalidated SVs were removed while those that failed in the validation assay design were subjected to manual review alongside SVs generated from four other studies (G4K, SCMC, ClinGen, and TARGET) in `BamViewer`^76^ and `Integrative Genomics Viewer` (`IGV`)^77^. To remove duplicate SVs with slightly different genomic coordinates — which may arise based on how SVs are detected at the two breakpoint locations — SVs were subjected to an automated curation process which refined the SV breakpoints by performing alignment of the reads at each breakpoint against a region defined by the breakpoint +/–200 bp using `BLASTN`^78^ v.2.6.0+.

In a recently published study^79^, we compared this approach with the consensus call approach, which requires an SV to be called by two algorithms (e.g. `SvABA`, `Manta`, and `Delly`). Using data from 560 Illumina-based WGS leukemia samples, we found that our curated SV data set missed 19% of multi-caller SVs while the multi-caller SVs missed about 2% of curated SVs that were either experimentally validated, had breakpoints matching focal deletion boundaries, or represented driver rearrangements consistent with the sample’s transcriptional subtype or RNA-detected fusions. The curated SVs missed by the multi-caller included *ETV6*-*RUNX1* rearrangements and reciprocals in 20 *ETV6*-*RUNX1* subtype samples, and structural variants representing focal *IKZF1* deletions in five patients. Given the extensive curation required for single-caller SVs, we opted to use the curated SV data set for this analysis.

632 TARGET samples were profiled by Complete Genomics Inc. (CGI). These SVs were obtained from our previous study^11^ in which SVs were identified using the CGI Cancer Sequencing service pipeline (version 2) and filtered to remove germline SVs. While performing the SV hotspot analysis, translocation hotspots joining *CHST13* and *GRK1*, as well as an inversion joining *GRIN2A* and the IGH locus, were found in multiple TARGET samples (AML, BALL, NBL, OS, and WT). As these hotspots were absent in samples from the same cancer type profiled by alternative sequencing technologies, they were considered PCR artifacts arising from the CGI sequencing approach, and we re-ran our analyses with the 97 variants linked to these hotspots excluded.

Duplicated samples from the same patient were consolidated by keeping the one with the highest number of SVs. As with the adult SVs, all pediatric variants were analyzed with the `FusionBuilder`^33^ pipeline, and entries identified as duplicates were removed.

### Copy number variation identification in pediatric cancer samples

`CONSERTING`^80^ v.1.0 was used to identify copy number changes in Illumina WGS samples after alignment to hg19 with `BWA`^75^ v.0.5.9. For CGI WGS data (including much of the TARGET data), we used copy number data from our previous analysis, which used an adapted version of `CONSERTING`^11^. Copy number profiles were manually corrected for tumor purity using allelic imbalance information for germline single-nucleotide polymorphisms.

### Chromothripsis analysis

Tumor samples characterized to have chromothripsis by previous studies^4,32,33,39,41,81–84^ were considered chromothripsis-positive. All samples were also analyzed by `ShatterSheek`^85^ v.1.1 using somatic CNVs and SVs to identify high-confidence chromothripsis events based on the following criteria: (1) at least six interleaved intra-chromosomal SVs, seven contiguous segments oscillating between two copy number states, the fragment joins test and either the chromosomal enrichment or the exponential distribution of breakpoints test or (2) at least three interleaved intra-chromosomal SVs and four or more inter-chromosomal SVs, seven contiguous segments oscillating between two copy number states and the fragment joins test. Candidates identified by `ShatterSeek` were subjected to manual inspection to finalize their chromothripsis status. *TP53* somatic and germline mutation status was acquired from the `ProteinPaint` data portal^86^.

### Complex genome rearrangement (CGR) analysis and the identification of complex SVs

Complex genome rearrangement (CGR) regions were identified using the default settings of `Starfish`^87^ v.0.11; in this context, `Starfish` defines CGR regions as complex structural changes that are likely to have originated in a single event. Once SVs within CGR regions had been identified, the remaining non-CGR SVs were analyzed using `ClusterSV` (https://github.com/cancerit/ClusterSV) to identify SVs that appeared to be part of additional complex events. SVs not identified with either application were considered simple SVs.

### Recombination signal sequence site identification

RSS sites were downloaded for the hg19 reference from the `RSSsite`^51^ website (https://www.itb.cnr.it/rss/). `RSSsite` assigns a quality score to each predicted RSS site in the form of a recombination information content (RIC) value; this value ranges from –1,000 (very poor) to 0 (very good) (https://www.itb.cnr.it/rss/help.html). There are two types of RSS sites with differing spacer lengths (12 or 23 bp), each of which has a different score threshold to be considered a potential site based on the cutoffs established by Cowell et. al (2002)^88^ (12 RSS: –38.81; 23 RSS: –58.45). All sites included in our analyses were required to exceed these threshold values. To aid in the visualization of RSS sites, we subtracted the threshold value from the RIC score for each site; thus, the RIC scores given in Supplementary Figures 9– 11 reflect the degree to which each site exceeds the threshold value, with higher values indicating sites with higher predicted recombination efficiencies. When multiple RSS sites overlapped, we merged the coordinate ranges for these sites and retained the best adjusted RIC score.

### Identification of SV breakpoint hotspots

In order to identify hotspots of SV breakpoints, we applied the nearest neighbor search to find the closest breakpoint pair across patient samples. We performed a comprehensive clustering analysis for SVs from the pediatric cancer cohort and the adult cancer cohort. Briefly, a breakpoint (BP) cut-off distance of 98 bp to the nearest neighbor (minimal distance between two BPs from different samples) was determined by `mclust`^89^ to identify two distinct clusters; breakpoints were then joined as segments if they had a nearest neighbor distance less than 98 bp.

The distance between identified breakpoint pairs serves as a parameter (*d*) for breakpoint segmentation across chromosomes. Breakpoints with *d* ≤ 98 bp in a window were joined as a segment (hotspot) as

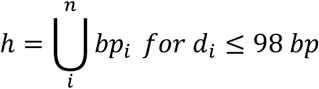

where *h*, *bp* and *n* indicate SV hotspot, breakpoint and patient sample size in the hotspot, respectively. For a given SV hotspot, *n* follows the Poisson distribution by

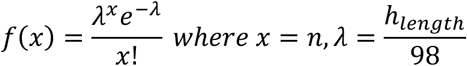

The *P-*value was calculated for each hotspot via the Poisson distribution and then subjected to multiple testing correction using the Benjamini and Hochberg method (FDR). Hotspots with FDR <0.05 are reported. Hotspots in the pediatric cohort required at least six breakpoints from multiple samples to exceed this threshold, while hotspots in the adult cohort required a minimum of eight.

### Classification of hotspots

Hotspots were classified as deletion (DEL), duplication (DUP), inversion (INV), or translocation (TRA) hotspots if at least 75% of the structural variants with a breakpoint at that hotspot were of the same type; otherwise, they were classified as other (OTH). The same approach was used to classify hotspots as originating from hematological malignancies, brain tumors, or solid tumors, with “multiple cancers” used for hotspots where no single category reached the 75% threshold. Hotspots were classified as “immune” if at least 75% of the breakpoints were within 1 Mb of the IGH, IGK, IGL, TRA, TRB, TRD, or TRG loci or were found within 1 Mb of one of the many IGH, IGK, IGL, or TRB orphan receptors located in the genome (such receptors are denoted as IGH-O, IGH-K, IGL-O, or TRB-O, respectively). Hotspots were also classified as “RSS” if at least 75% of the breakpoints were within 20 bp of an RSS site predicted by `RSSsite`.

### SV signature analysis

To ensure the quality of the SV signature analysis, we excluded 65 tumors with spurious copy number changes associated with over-segmentation (Supplementary Table 1). Spurious copy number profiles were identified by visually inspecting the copy number profile of each sample to find those with an abundance of small and subclonal copy number losses or gains accompanied by a lack of SV support. Such events gave the appearance of a “fractured genome” and were caused by library construction artifacts as documented in our prior study^80^.

`SigProfilerMatrixGenerator`^90^ v.1.2.29 was used to classify each somatic SV into one of 32 SV types as done previously^19,21^, based on the SV type, the distance between the SV breakpoints (for intra-chromosomal SVs; SVs under 1 kb in size were filtered out), and whether or not the SV was found to cluster with other SVs within the same sample. SV signatures were then extracted from 343 pediatric cancer samples and 1,656 adult samples, each at least 20 SVs (to increase signature robustness by ignoring low-burden samples which add noise), using `SigProfilerExtractor`^52,53^ v.1.1.24. One to 15 signatures were extracted from each dataset using the “double” precision mode and 500 NMF replicates per signature. Based on stability and reconstruction error scores, `SigProfilerExtractor` determined the ideal number of signatures to be eight for the pediatric samples (Supplementary Figure 22) and ten for the adult samples (Supplementary Figure 23). The abundance of these eight and ten signatures was measured across all pediatric and adult cancers, respectively, using `SigProfilerAssignment`^91^ v.0.1.8. To determine the percentage of cancers bearing each SV signature, we considered a sample positive for a given signature if at least one SV was assigned to the signature in the sample. For samples with a reconstruction cosine score <0.80 (comparing the actual SV profile with the reconstructed profile), all signatures were set to zero (non-detected) for this analysis to remove false positives while keeping the sample in the denominator.

All eight pediatric and ten adult SV signatures were compared to the ten SV signatures reported in the COSMIC v.3.4 database (https://cancer.sanger.ac.uk/signatures/sv/) by `SigProfilerAssignment`^91^ v.0.1.8. Reconstruction cosine scores had to be ≥0.80 for a signature to be considered a match.

### SV evolution analysis

For multi-sample SV evolution analysis, SVs were detected using `CREST`^74^ v.1.0 as described above. SVs were clustered by their presence/absence within each sample for each patient. As the same SV may have a slightly different call in two different samples (such as genomic coordinates that vary by a few bp or a miss due to a low read count), we ran a search using the SV junction sequence against the raw NGS reads to determine the read count in all samples from the same patient for each SV.

### Motif analysis

*De novo* motif analysis was performed using the default parameters of `HOMER`^49^ v.4.11. DNA sequences from SV hotspots were appended 20 bp to each side (left and right).

### HRD features

Curated somatic SNV data was downloaded from the St. Jude Cloud Pediatric Cancer Knowledgebase (PeCan)^92^. Small indels were called by `Mutect2`^93^ from `GATK` v4.1.8.0 without quality filtering, and artifacts due to mapping and germline contamination were filtered out by running `indelPost`^94^ to remove matches to `gnomAD`^95^ v2.1.1. Those, together with curated SVs, were used to run `CHORD`^55^ for 976 samples profiled by Illumina sequencing, as samples profiled by CGI had a high rate of PCR artifacts, which can compromise indel re-analysis. HRD detection was performed using a random forest model.

### Analysis of non-coding regulatory SVs leading to oncogene cis activation

We focused on well-characterized non-coding drivers in relevant cancer types to summarize the prevalence of such events. The criteria used for each region is described as follows:

*MYC* regulatory regions were identified for B-ALL, T-ALL and neuroblastoma. These included the NOTCH-MYC enhancer (NME)^96^, blood enhancer cluster (BENC)^97^ and PVT1^98^ and known enhancer hijacking events in neuroblastoma^99^. The genomic regions for NME and BENC were sourced from Lancho and Herranz (2018)^100^ and converted to the hg19 coordinates used in this study (chr. 8: 130,161,775– 130,202,012 for NME and chr. 8: 130,558,973–130,724,424 for BENC). The genomic region upstream of MYC and the end of PVT1 (hg19: 128,753,680–129,113,499) was used to query events that disrupted the PVT1 and MYC regulatory region. NME and BENC regions were queried for duplication SVs while the PVT region was queried for translocation events and any events that could disrupt PVT transcription.

For deletions upstream of *TAL1* in T-ALL, those with breakpoints either within a portion of *TAL1* (chr. 1: 47,681,962–47,698,007; reverse) or upstream, including within all or a portion of *STIL* (chr. 1: 47,715,811– 47,779,819; reverse) were considered non-coding drivers.

*DUX4* is activated by translocation to *IGH* locus in B-ALL, which involves highly repetitive regions at both loci^28,101^. In both research and clinical settings, such events are primarily detected by overexpression of *DUX4* in RNA-seq or cytogenetics profiling. We were able to find WGS SV breakpoints for eight of these, which is likely an underestimate.

*TERT* activation in neuroblastoma^102^ was identified based on inter-or intra-chromosomal translocations with breakpoints either upstream of *TERT* (chr. 5: 1,925,184–1,325,183) or 6–40 Kb downstream. Deletions or amplifications encompassing the genomic region encoding *TERT* gene (chr. 5: 1,253,282–1,295,183; transcribed in reverse orientation) were excluded.

*GFI1* and *GFI1B* activation in medulloblastoma^103^ was driven by inter-and intra-chromosomal rearrangements, respectively, based on prior publications. For *GFI1*, translocations with a breakpoint within 102 Kb upstream or downstream of *GFI1* were selected. For GFI1B, intra-chromosomal events were selected if they minimally overlapped with at least a portion of chr. 9: 134,279,237–1,358,562,045 (determined from Northcott et al. (2014)^103^) and contained breakpoints upstream or within *GFI1B*. Deletion and duplication events were required to be <4 Mb size and those that overlapped the entire *GFI1B* gene were excluded.

*BCL11B* activation in leukemia^104^ was identified based on duplications in the region 730 Kb downstream as well as breakpoints within 100 Kb downstream and upstream of *BCL11B* with breakpoint partners in the gene desert upstream of *ARID1B* on chr. 6, the blood enhancer cluster (BENC) distal to *MYC* within the *CCDC26* gene (chr. 8), *CDK6* (chr. 7), *ETV6* (chr. 12), or *SATB1* (chr. 3).

SNCAIP promoting *PRDM6* activation in medulloblastoma^105^ was based on intra-chromosomal events that overlap *SNCAIP* and were no larger than 1 Mb in size.

*FLT3* activation in ALL^69^ was based on deletions with at least one breakpoint upstream of *FLT3* within the coordinate range of chr. 13: 28.5–28.9 Mb, and with a total span of <1 Mb.

*MECOM* in AML^106^ was based on events that had breakpoints within +/–100 Kb of *MECOM*, except for those events with breakpoints within the gene or that completely duplicated or deleted the gene.

*CEBPD* in ALL^107^ was based on translocation events with breakpoints within +/–100 Kb of *CEBPD*.

*CRLF*2 in ALL^67^ was based on events with breakpoints within +/–100 Kb of *CRLF2* except for deletions which deleted part of the gene itself.

## Abbreviations

Pediatric cancer types are abbreviated as follows

ACT: adrenocortical cancer
AML: acute myeloid leukemia
B-ALL: B-lineage acute lymphoblastic leukemia
BT other: brain tumor, other
EPD: ependymoma
EWS: Ewing sarcoma
HGG: high-grade glioma
HM other: hematological malignancy, other
LGG: low-grade glioma
MB: medulloblastoma
MEL: melanoma
NBL: neuroblastoma
OS: osteosarcoma
RB: retinoblastoma
RHB: rhabdomyosarcoma
RT: rhabdoid tumor
ST other: solid tumor, other
T-ALL: T-lineage acute lymphoblastic leukemia
WT: Wilms tumor

## Supplementary figures and legends

**Supplementary Figure 1.**
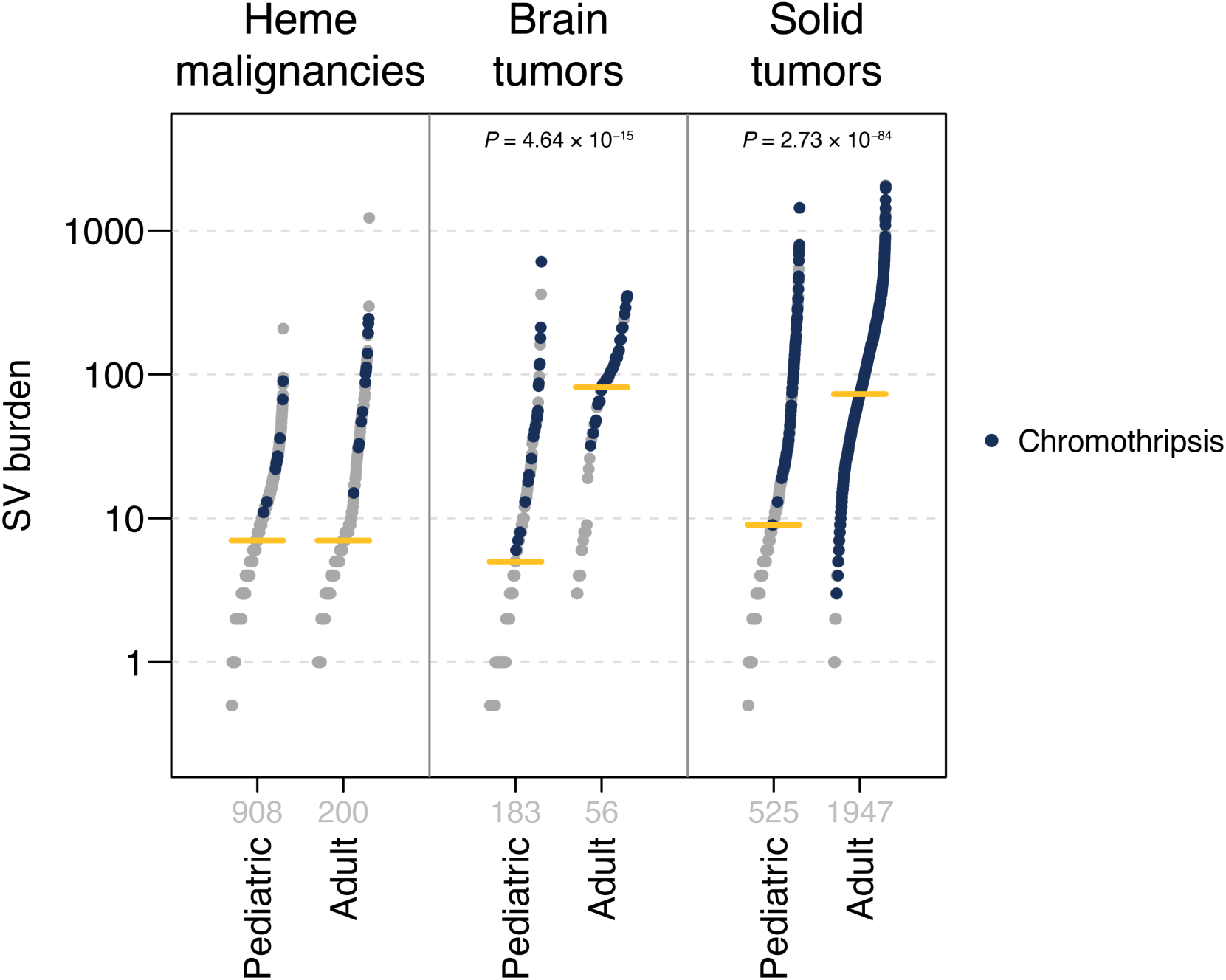
Pediatric and adult SV burden by cancer category. Plots show the SV burden of pediatric (left) and adult (right) samples in hematological malignancies, brain tumors and solid tumors. Each point represents one sample, blue dots indicate samples with chromothripsis, and median values are indicated with yellow lines. Sample numbers for each cancer type are listed in gray. Significant *P*-values by two-sided Wilcoxon rank-sum tests are listed for categories with significant differences in the SV burden between pediatric and adult samples.

**Supplementary Figure 2.**
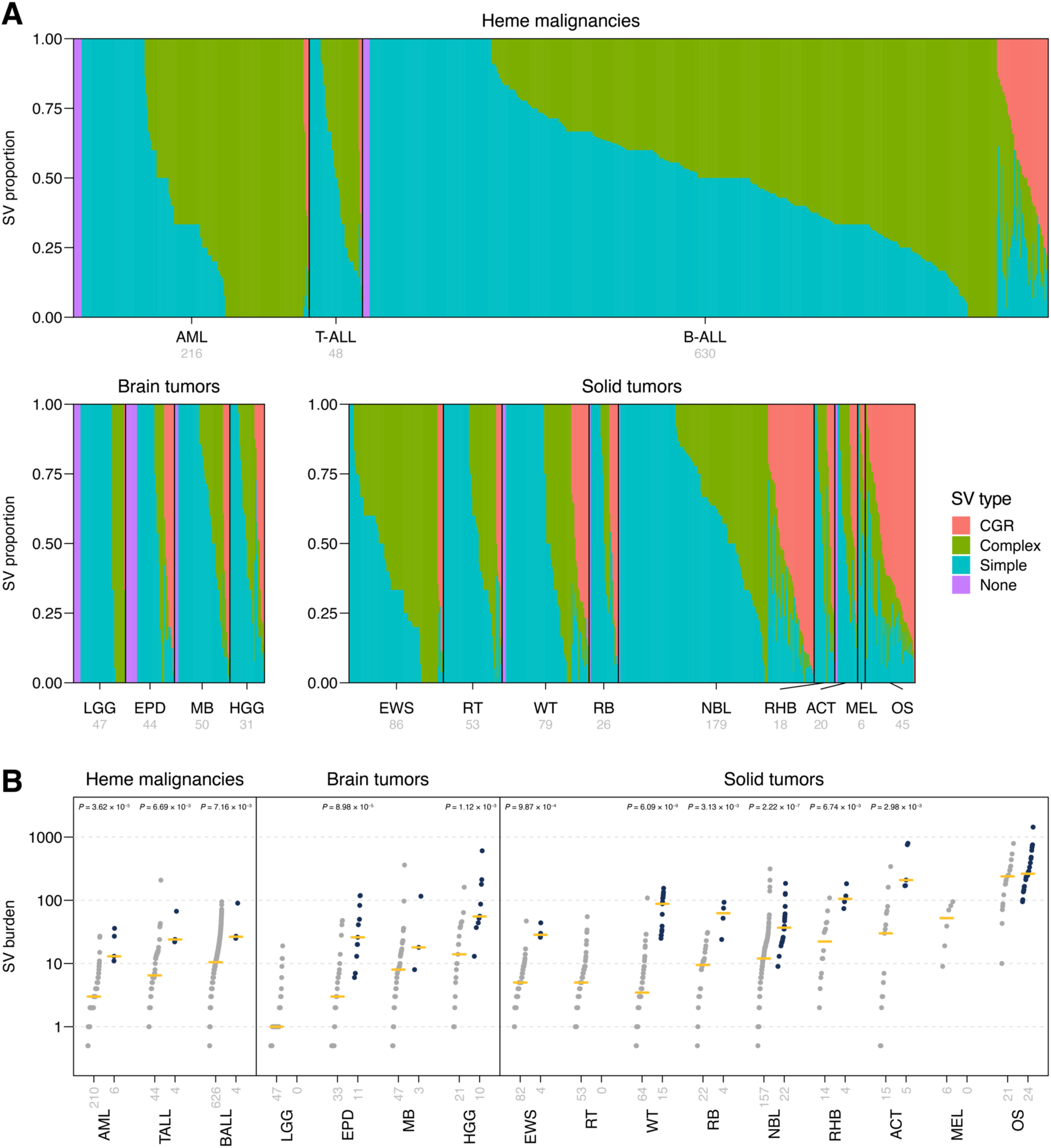
Identification of complex SVs and chromothripsis in pediatric cancers. (**a**) Stacked bar chart showing the proportion of structural variants classified as part of complex genome rearrangements (CGR, pink) likely originating from a single event, complex events (green) which refer to additional clustered SVs, or simple events (blue) in each sample. Bars for samples with no detected SVs are shown in magenta. Sample numbers for each cancer type shown in gray. (**b**) SV burden of chromothripsis-negative (left, gray) and chromothripsis-positive (right, blue) samples, divided by cancer category. Each point represents one sample, and median values are indicated with yellow lines. Sample numbers for each cancer type are listed in gray. Significant *P*-values by two-sided Wilcoxon rank-sum tests are listed for cancers with significant differences in the SV burden between chromothripsis-negative and chromothripsis-positive samples.

**Supplementary Figure 3.**
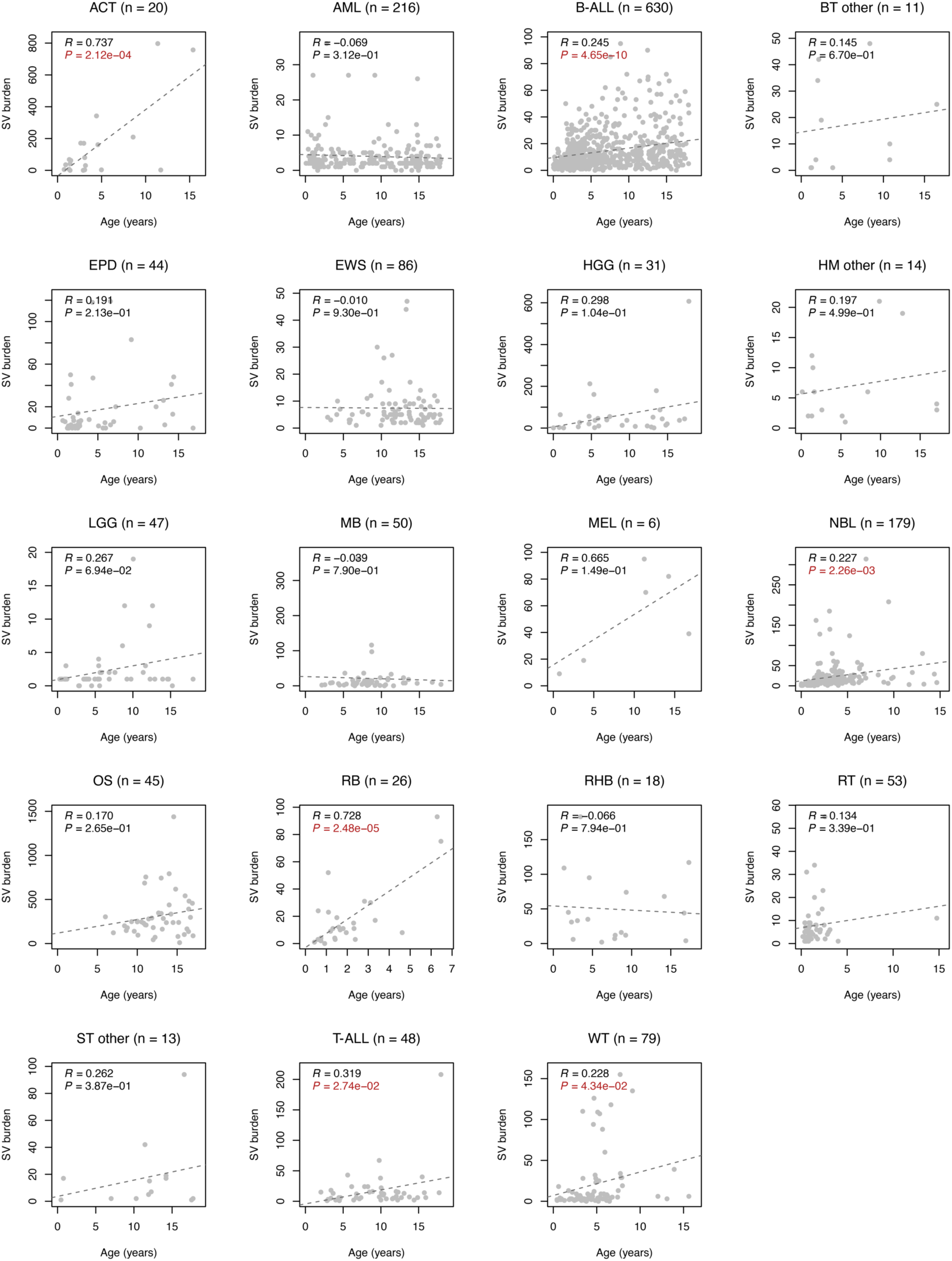
SV burden by age in pediatric cancers. Each plot shows one pediatric cancer type, with x-axis indicating age at diagnosis and y-axis the total number of SVs in a sample. Each point represents one sample, and sample numbers for each cancer type are shown at top. Pearson *R* and *P*-values are also shown, with significant *P*-values shown in red.

**Supplementary Figure 4.**
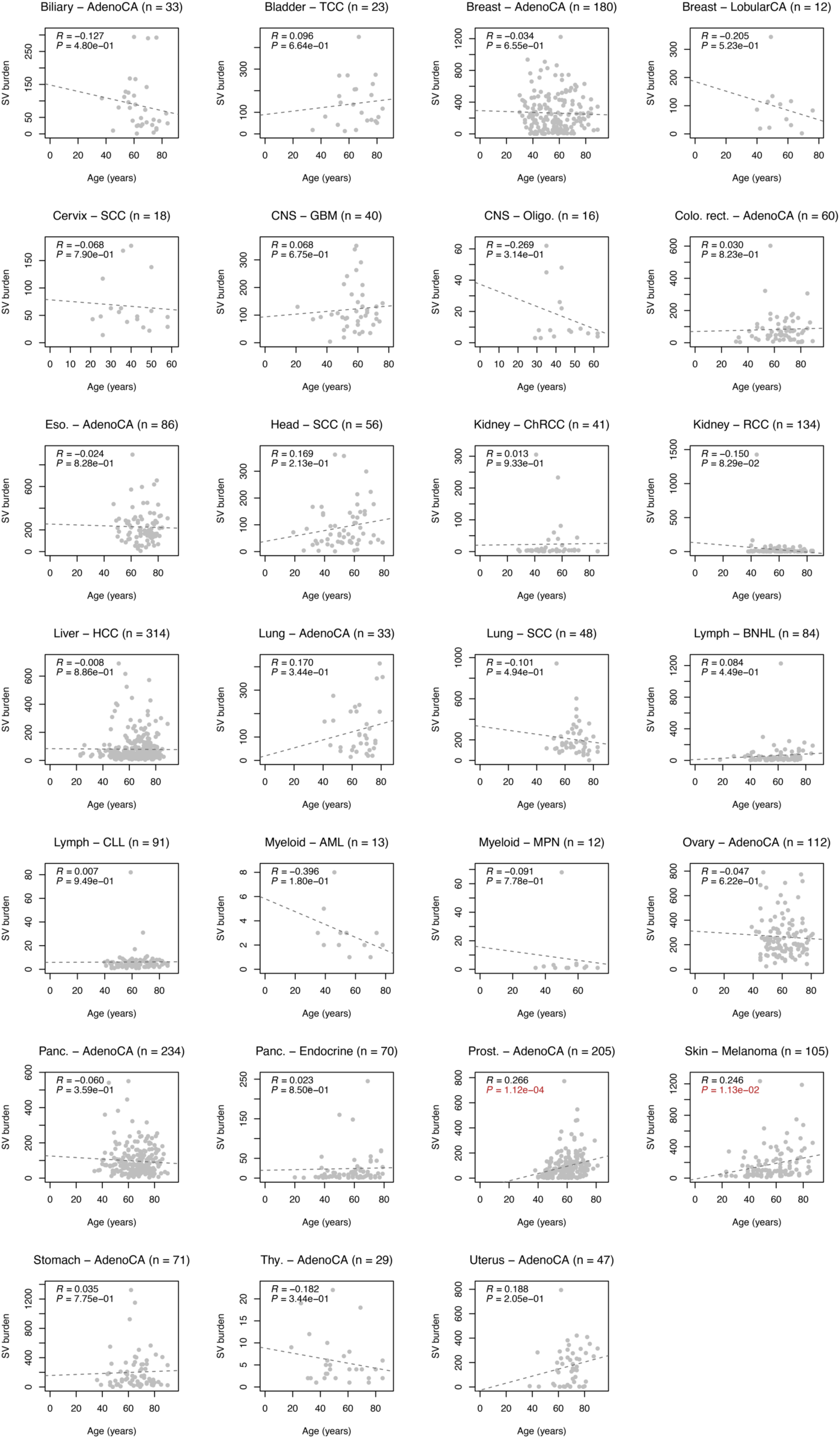
SV burden by age in adult cancers. Each plot shows one adult cancer type (using PCAWG data), with x-axis indicating age at diagnosis and y-axis the total number of SVs in a sample. Each point represents one sample, and sample numbers for each cancer type are shown at top. Pearson *R* and *P*-values are also shown, with significant *P*-values shown in red.

**Supplementary Figure 5.**
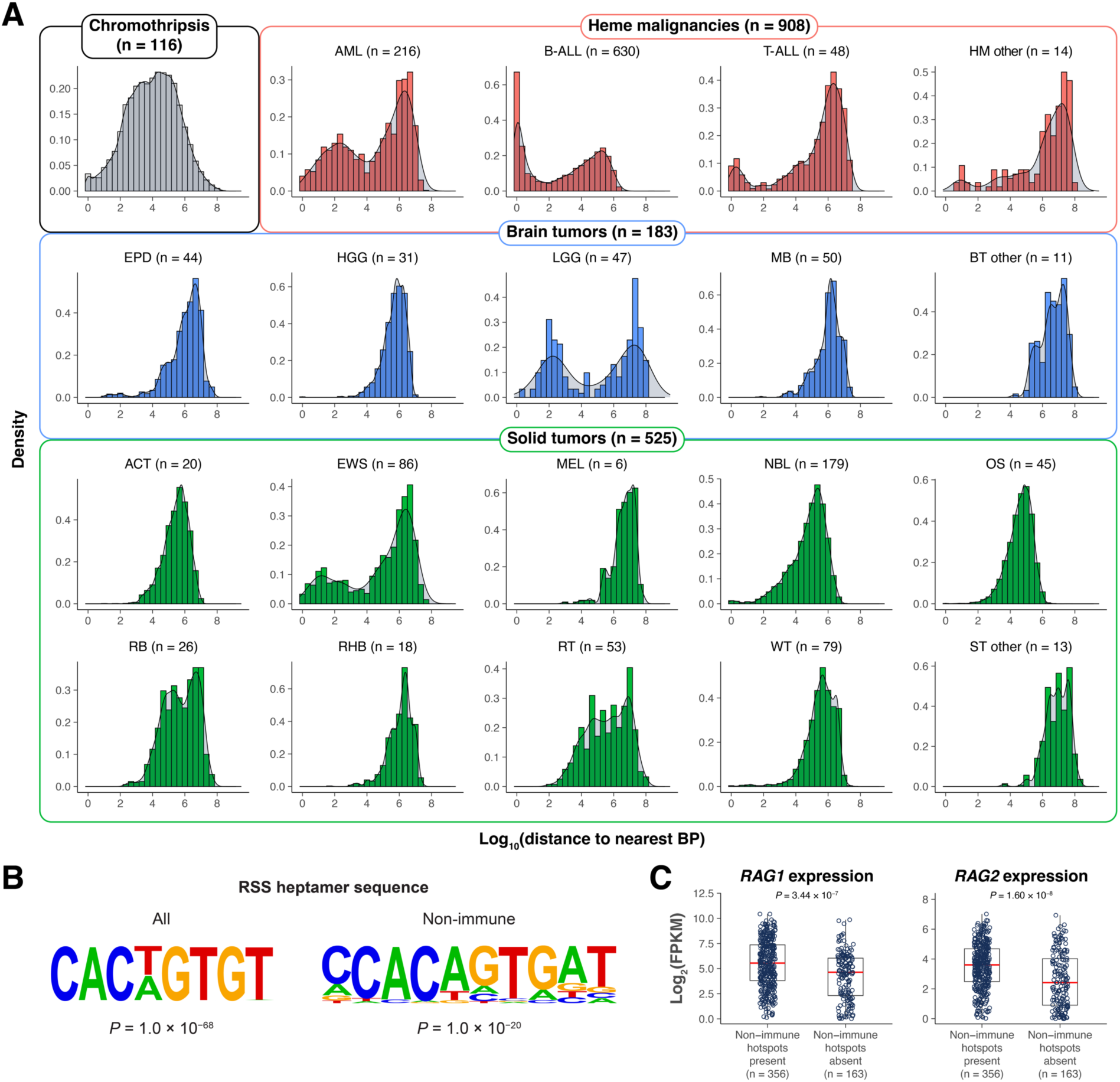
Somatic SV hotspots in pediatric cancer. (**a**) Distribution of distance to the nearest breakpoint (BP) within chromothripsis-positive samples and in each pediatric cancer type. (**b**) Enrichment of RSS heptamer sequence in all hotspots and non-immune hotspots. (**c**) *RAG1* and *RAG2* expression in B-ALL samples with and without non-immune SV hotspots. *P*-values are obtained via two-sided Wilcoxon rank-sum test.

**Supplementary Figure 6.**
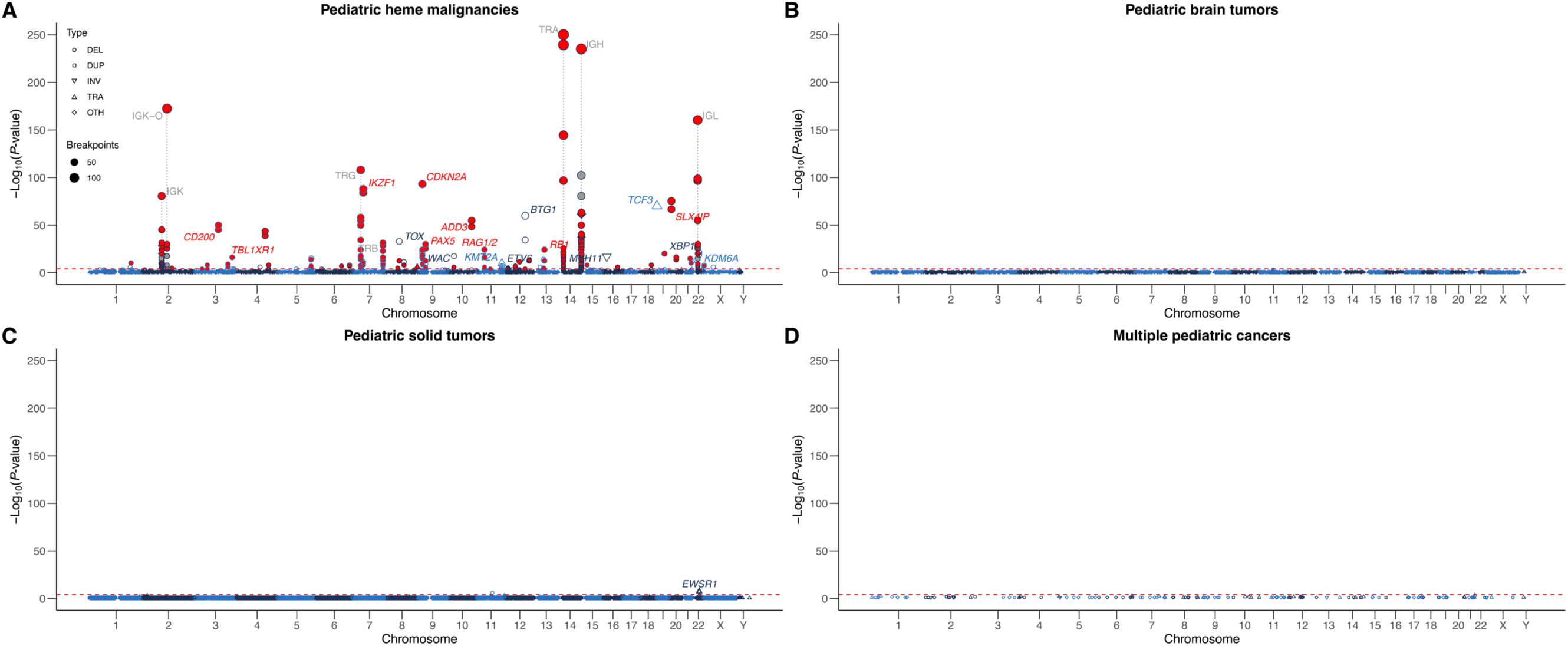
Somatic SV hotspots of pediatric cancer. Manhattan plots showing SV hotspots contributed by pediatric (**a**) hematological malignancies, (**b**) brain tumors, (**c**) solid tumors, and (**d**) multiple cancer types, and their significance across chromosomes is shown in the same style as Fig. 2D. Hotspots were classified as contributed by heme malignancies, brain tumors, or solid tumors if at least 75% of the breakpoints were contributed by cancers within that category; otherwise, they were assigned to the category of “multiple pediatric cancers”.

**Supplementary Figure 7.**
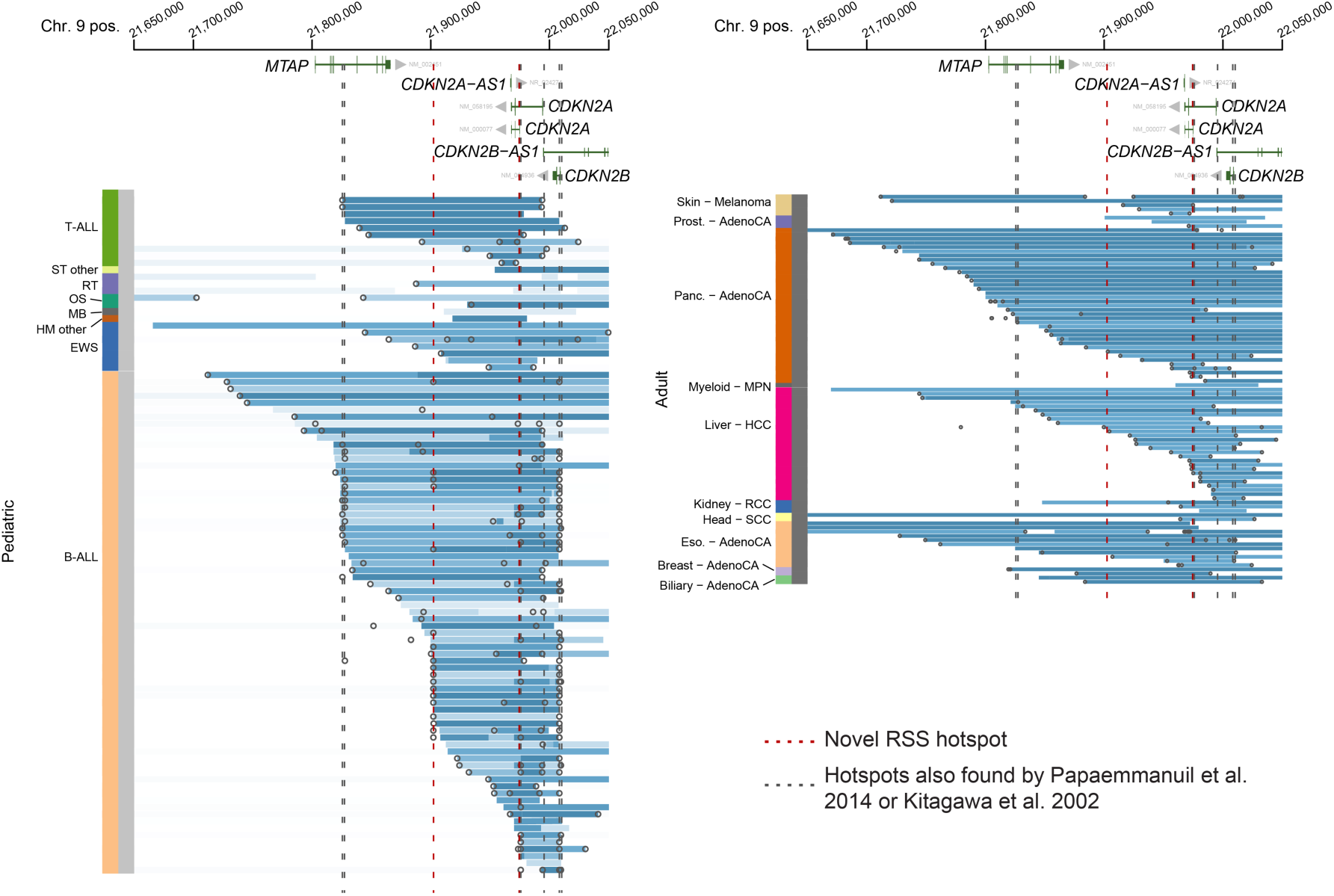
Somatic SV hotspots in *CDKN2A*. Copy number and SV profiles for pediatric (left) and adult (right) cancers with *CDKN2A* deletions. Each row represents one sample, with cancer type indicated on the left. Circles denote the locations of SV breakpoints while blue coloring indicates copy loss. Vertical dotted lines indicate the locations of RSS associated with SV hotspots in B-ALL found in this study (red color) or reported in previous studies (gray color).

**Supplementary Figure 8.**
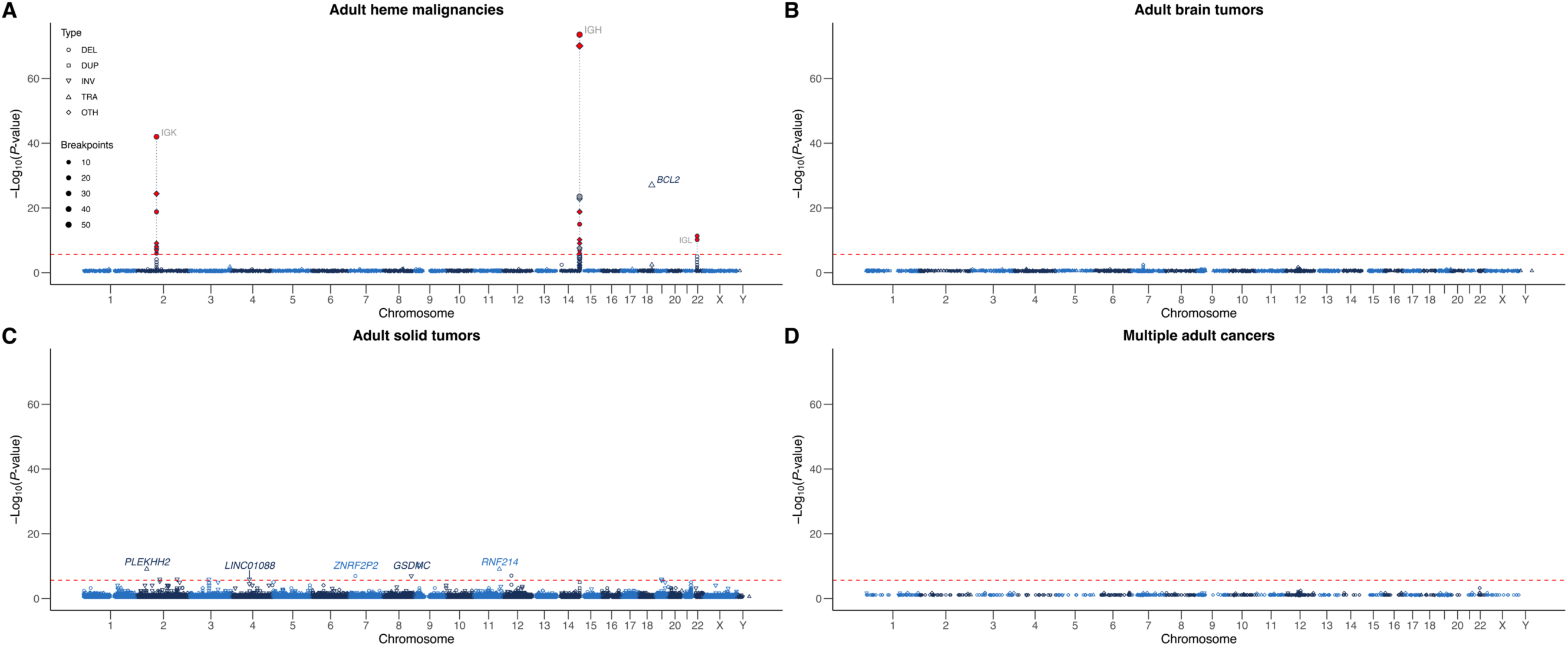
Somatic SV hotspots of adult cancer. Manhattan plots showing SV hotspots contributed by adult (**a**) heme malignancies, (**b**) brain tumors, (**c**) solid tumors, and (**d**) multiple cancer types, and their significance across chromosomes is presented in the same style as Supplementary Figure 6.

**Supplementary Figure 9.**
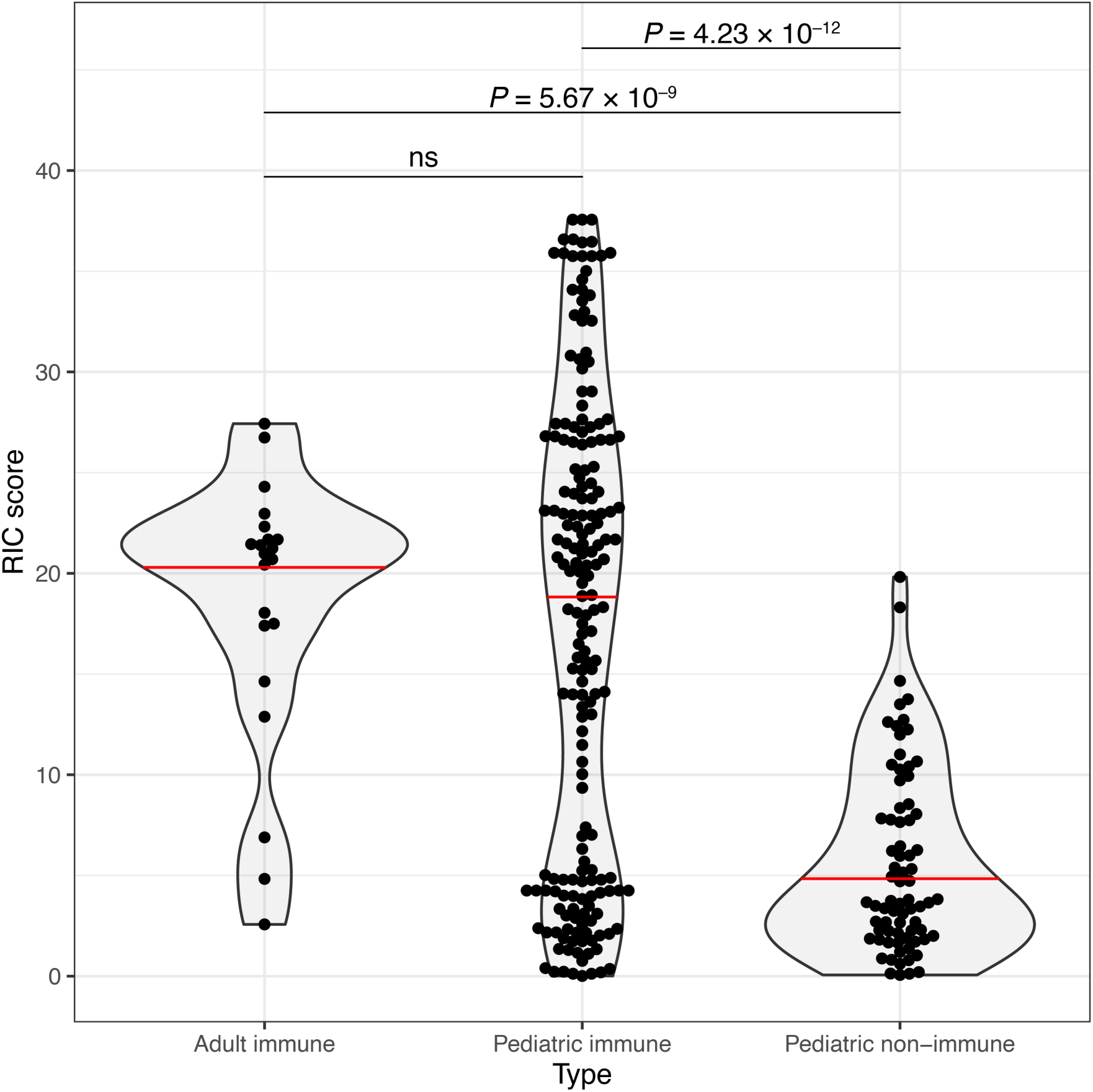
Distribution of recombination information content (RIC) scores of predicted RSS sites in immune and non-immune hotspots in pediatric and adult cancers. Violin plots of RIC scores associated with hotspots in adult immune regions, and pediatric immune and non-immune regions. For ease of visualization, RIC scores are presented as the score above the threshold value required for positive identification by `RSSsite` (see Methods). Red lines indicate group medians. Significant *P*-values by two-sided Wilcoxon rank-sum tests are listed for each comparison and are corrected for multiple testing using the Benjamini and Hochberg method.

**Supplementary Figure 10.**
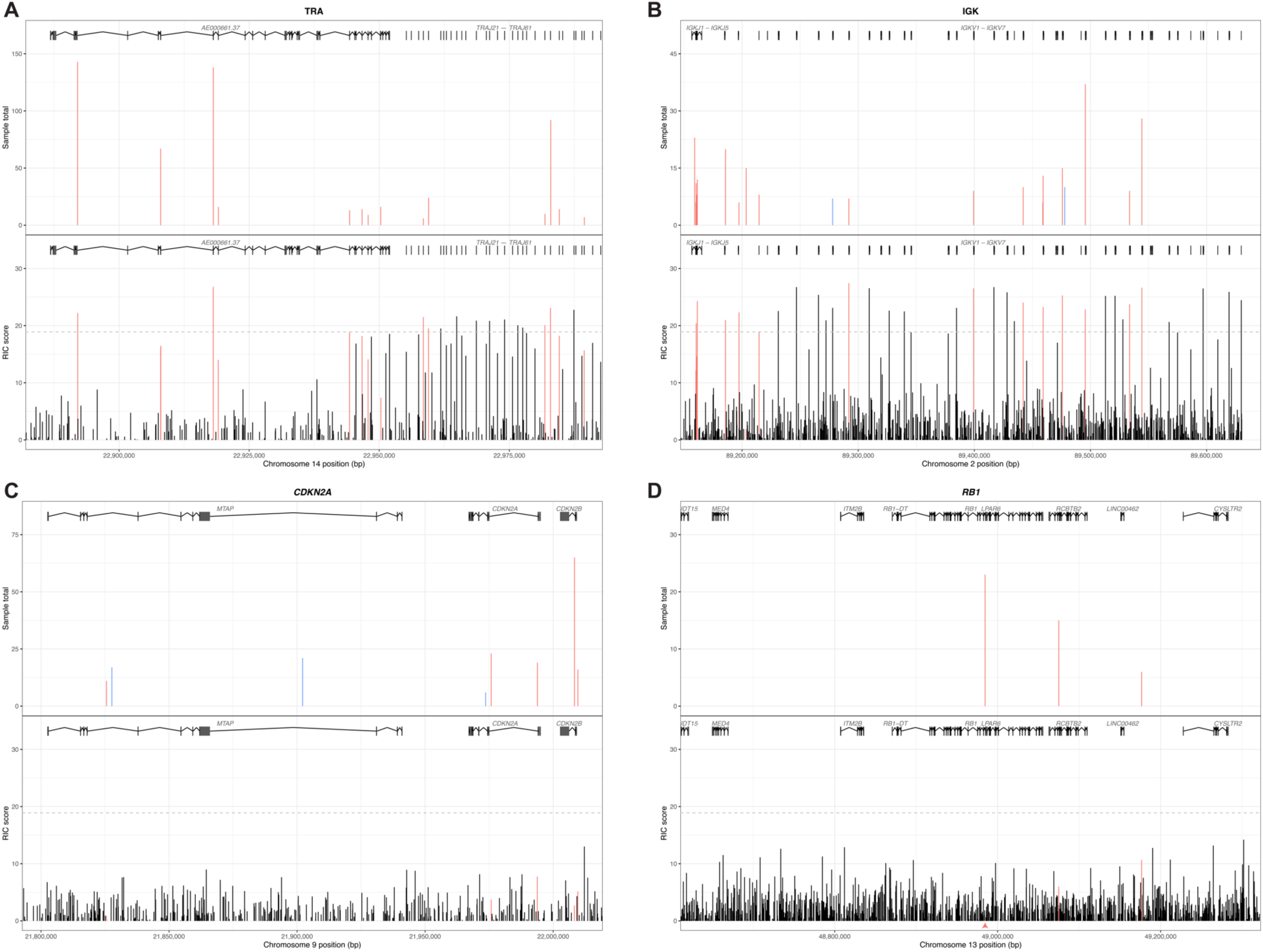
Differential RIC scores of pediatric SV hotspots in representative immune and non-immune regions. SV hotspots in immune regions are shown in (**a**) TRA and (**b**) IGK while those in non-immune loci are shown in (**c**) *CDKN2A* and (**d**) *RB1*. Gene models are displayed at the top with the x-axis indicating the genomic position. Top panels show the SV hotspots with the total number of samples with SV breakpoints shown on the y-axis and colored by their RSS association status (red: RSS-associated; blue: not RSS-associated). Bottom panels show the distribution of all predicted RSS sites, with the recombination information content (RIC) score given on the y-axis; for ease of visualization, RIC scores are presented as the score above the threshold value required for positive identification by `RSSsite` (see Methods). RSS sites are colored red if associated with a SV hotspot in the top panel. The dotted line indicates the median RIC score for RSS sites associated with pediatric immune region hotspots (18.9). The red arrowhead in (d) denotes a low-scoring site (0.2 above the minimum threshold RIC score) that is linked to SV breakpoints from 23 samples (i.e. RAG RSS #1 site in Figure 2).

**Supplementary Figure 11.**
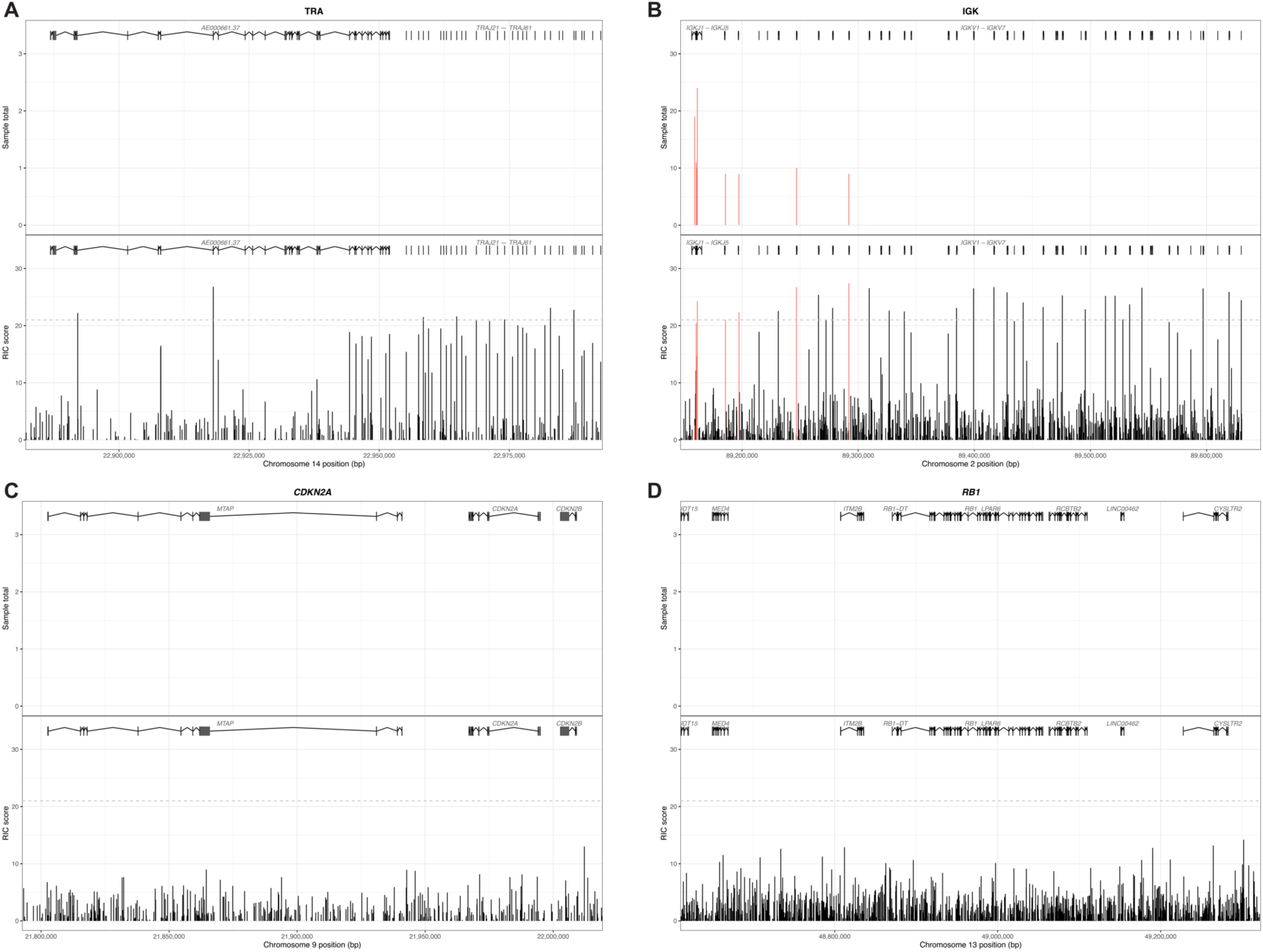
RIC scores of adult SV hotspots in representative immune and non-immune regions. The SV hotspots were based on adult PCAWG data in the same four regions as those shown for the pediatric SV hotspots in Supplementary Figure 10 using the same style. SV hotspots were detected in IGK (**b**) but absent in TRA (**a**), CDKN2A (**c**), and RB1 (**d**). The dotted line indicates the median RIC score for RSS sites associated with adult immune region hotspots (21.0).

**Supplementary Figure 12.**
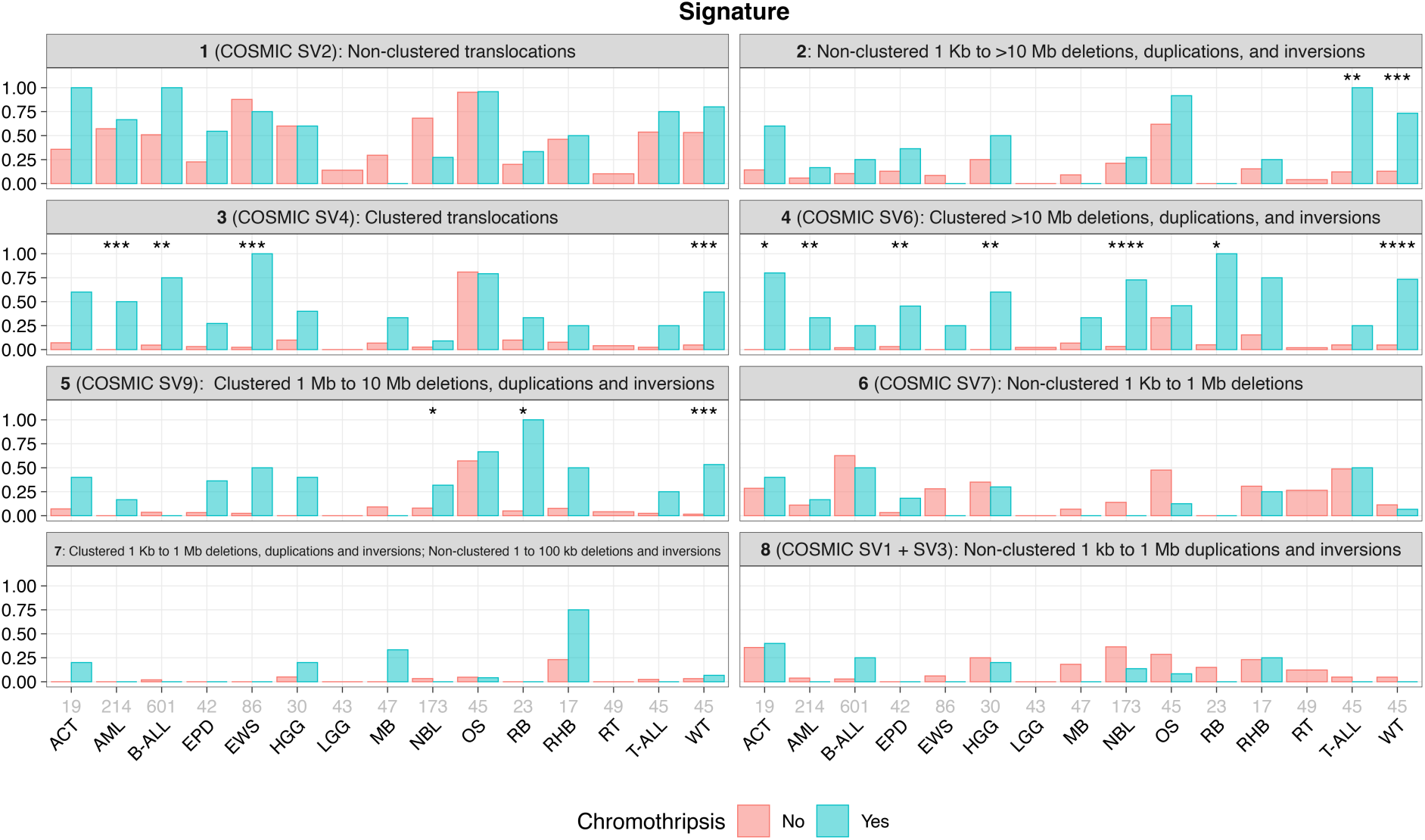
SV signature associations with chromothripsis. Proportion of chromothripsis-negative (red bar) and chromothripsis-positive samples (blue bar) for all cancer types with at least 15 samples in the pediatric cohort exhibiting each of the eight signatures. Matching COSMIC signatures (if present) and a brief description of each signature are provided. Significant *P*-values by Fisher’s exact test are noted for each signature and cancer type with *, **, ***, and ****, which represent *P*-values ≤0.05, 0.01, 0.001, and 0.0001, respectively.

**Supplementary Figure S13.**
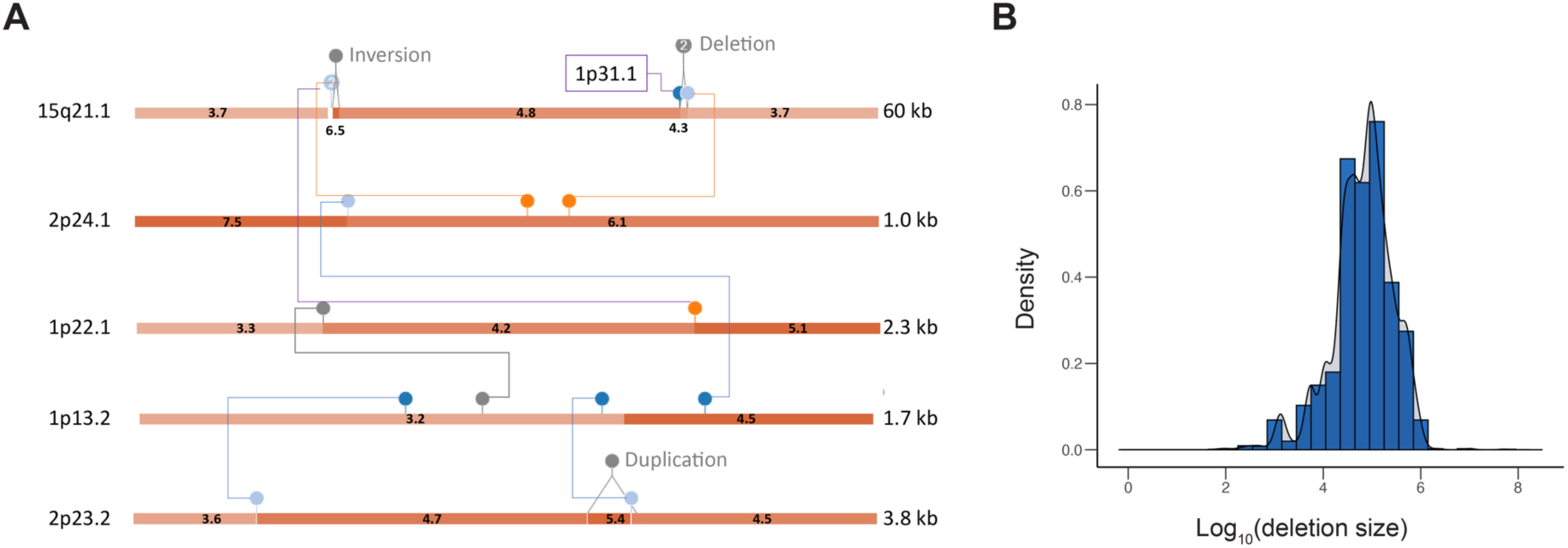
Illustration of features representative of pediatric SV signatures 3 and 6. (**a**) An example of clustered inter-chromosomal events in SJOS013, the defining feature of pediatric SV signature 3. This event involves five distinct regions of chromosomes 1, 2, and 15 contain 2–4 rearrangements within 1–2 kb. SVs defined as inter-chromosomal translocations are shown with colored dots while intra-chromosomal events are shown with gray dots. (**b**) Histogram of deletion sizes associated with RSS hotspots. The deletion sizes match the non-clustered deletions comprising pediatric SV signature 6 (i.e. non-clustered deletion between 1 Kb – 1 Mb), which is enriched in pediatric B-ALL and T-ALL.

**Figure S14.**
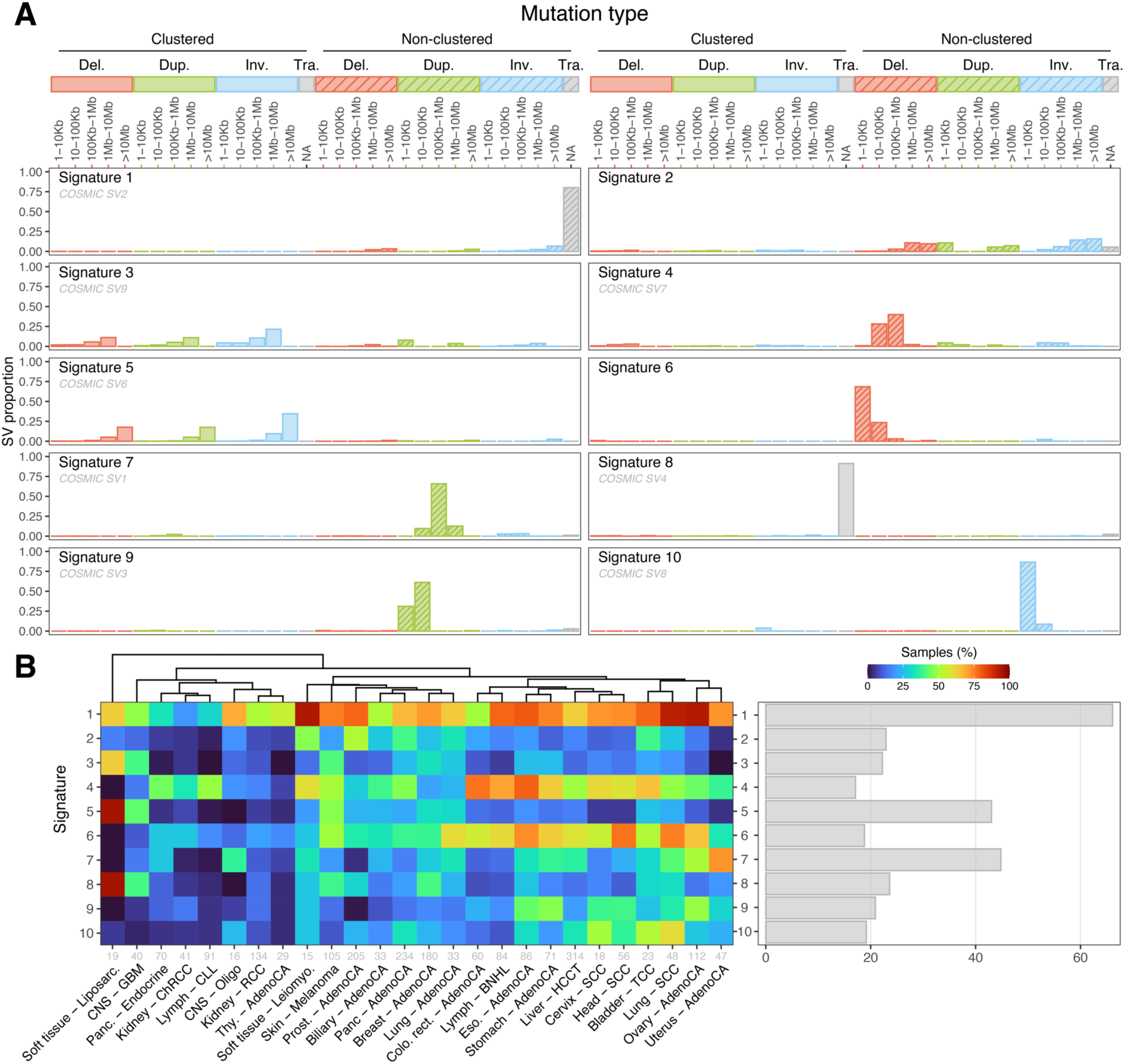
Somatic SV signatures in adult cancer. (**a**) Bar plots showing the distributions of the 32 SV types in all ten SV signatures extracted from the adult cancer cohort. Non-clustered signatures are represented by patterned bars. Signatures with matches to the COSMIC database have their corresponding COSMIC signature labeled in gray. (**b**) Heatmap showing the percent of cancer samples with each of the SV signatures shown in (a) for all cancer types with at least 15 samples. The bar plot at right shows the percentage of all adult cancer samples exhibiting each signature.

**Supplementary Figure 15.**
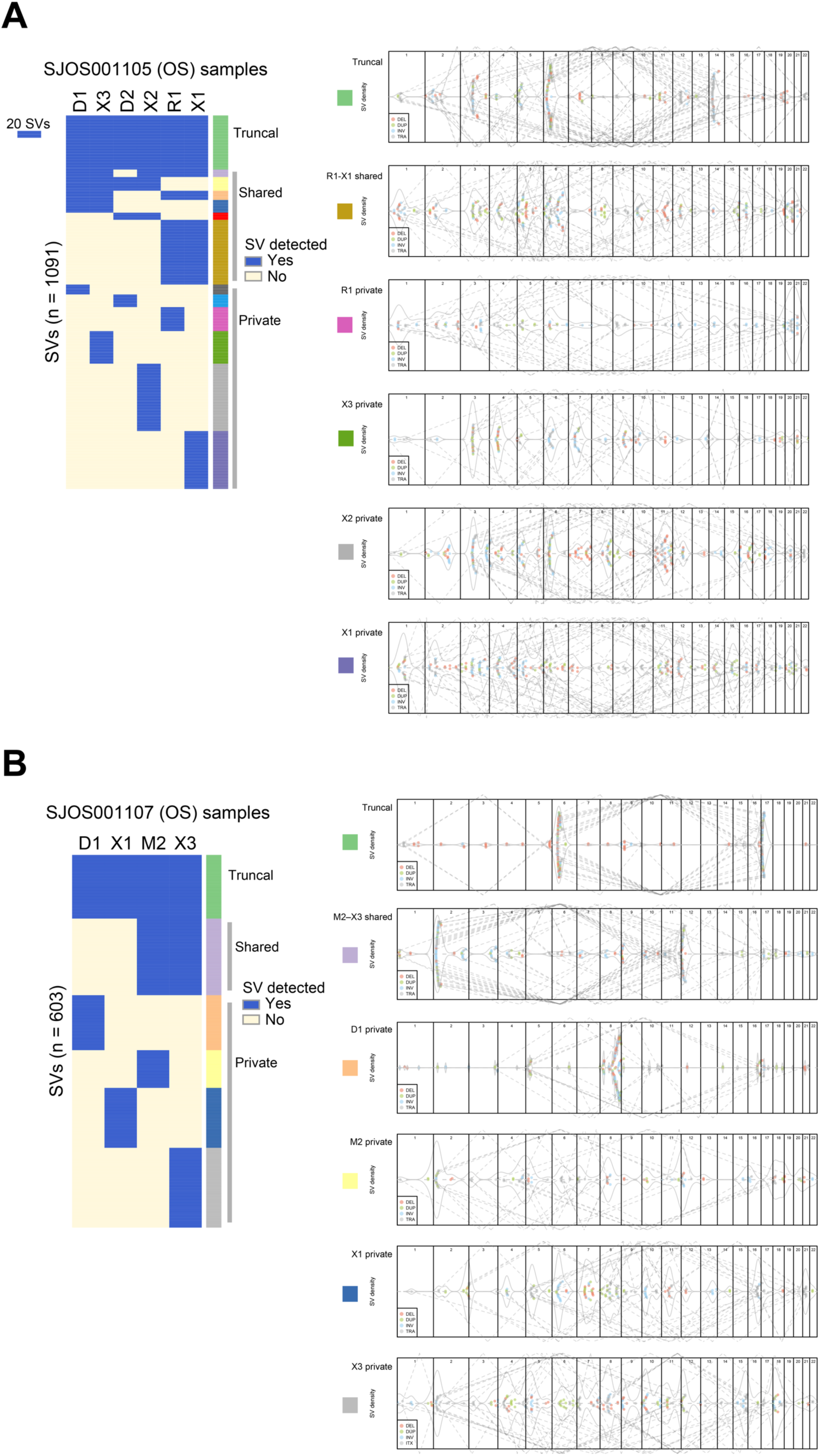
SV temporal evolution in two multi-sample osteosarcoma patients. SV grouping (left) and genome-wide density plot (right) shown in the same style as Figure 4. (**a**) Diagnostic (D1) and metastatic (D2 and R1) tumors profiled for patient SJOS001105 with D2 and R1 samples acquired two years after D1. X3, X1 and X2 are from PDX models developed from D1, R1 and M2, respectively.(**b**) Diagnostic (D1) and metastatic (M2) tumors profiled for patient SJOS001107, with the M2 sample acquired one year after D1. X1 and X3 are from PDX models developed from D1 and M2, respectively. In both patients, presence of SVs private to PDX samples indicates that PDX samples appear to grow out of one of the clones of the primary patient tumor samples and may have continued SV evolution.

**Supplementary Figure 16.**
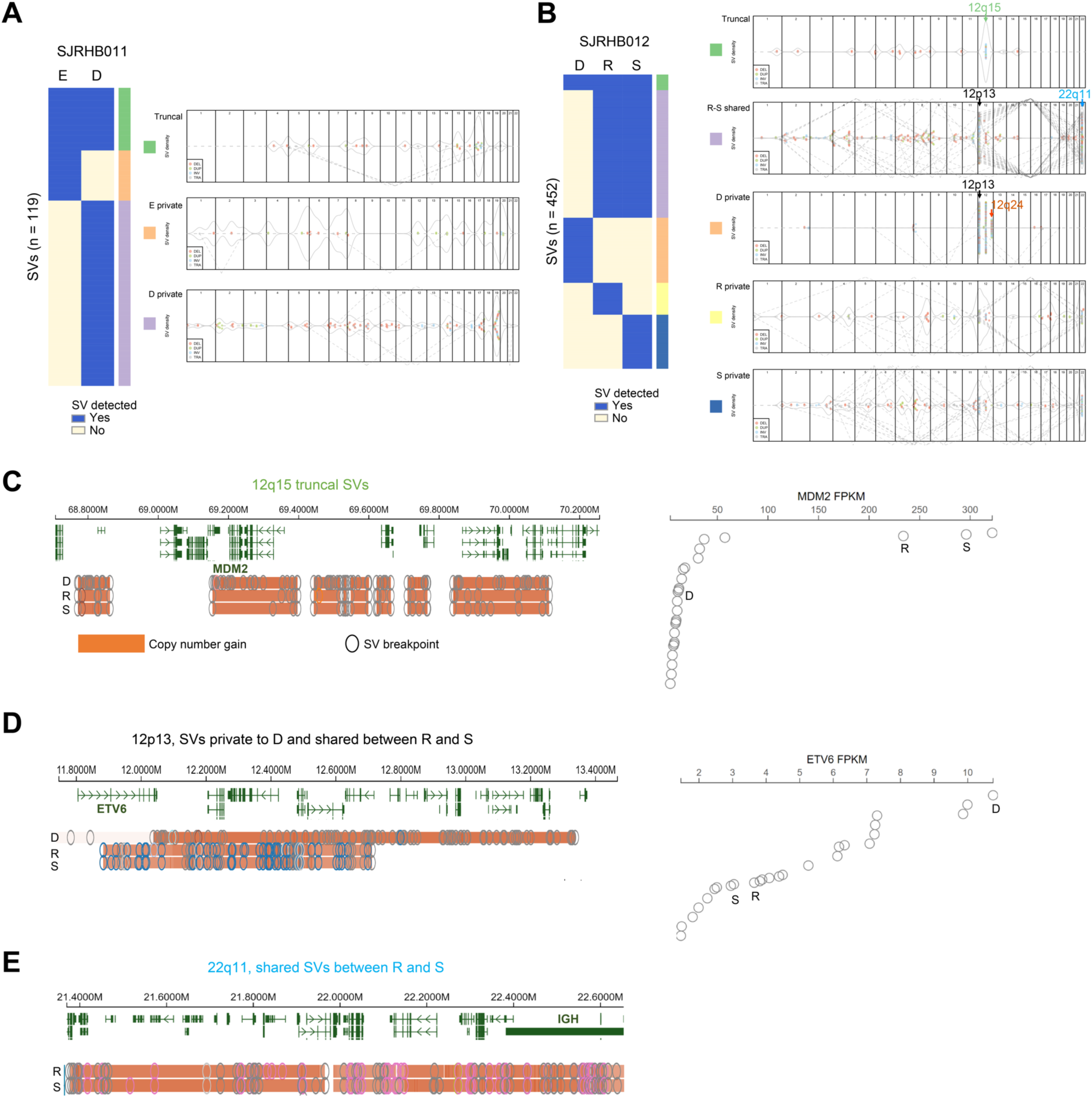
SV evolution in two embryonal rhabdomyosarcoma (ERMS) patients. (**a**) SV grouping (left) and genome-wide density plot (right) shown in the same style as Figure 4 for two RHB samples obtained from one patient. In this particular case, E refers to tumor sample acquired at diagnosis and D refers to a relapsed tumor acquired 15 months later. (**b**) As in (a), except that a different patient is shown. In this case, D was from diagnosis at prostate while R and S were from relapsed tumors at prostate and pelvis acquired 14 months later. Chromosome 12 had three different groups of SVs: 12q15 with truncal variants (shown in **c**), 12p13 which can form either a shared SV group joining 22q11 (details in **d** and **e**) or a private SV group joining 12q23. (**c**) Truncal copy-number gain (left) leading to increased *MDM2* expression (right), particularly in R and S in sample SJRB012 shown in (b). The gene expression data were based on all ERMS samples hosted on the `GenomePaint` portal (https://viz.stjude.cloud/tools/genomepaint). (**d**) Private SVs leading to increased *ETV6* expression in D; shared SVs impacting R and S are present in the same region. (**e**) Shared SVs between R and S impacting the *IGH* locus.

**Supplementary Figure 17.**
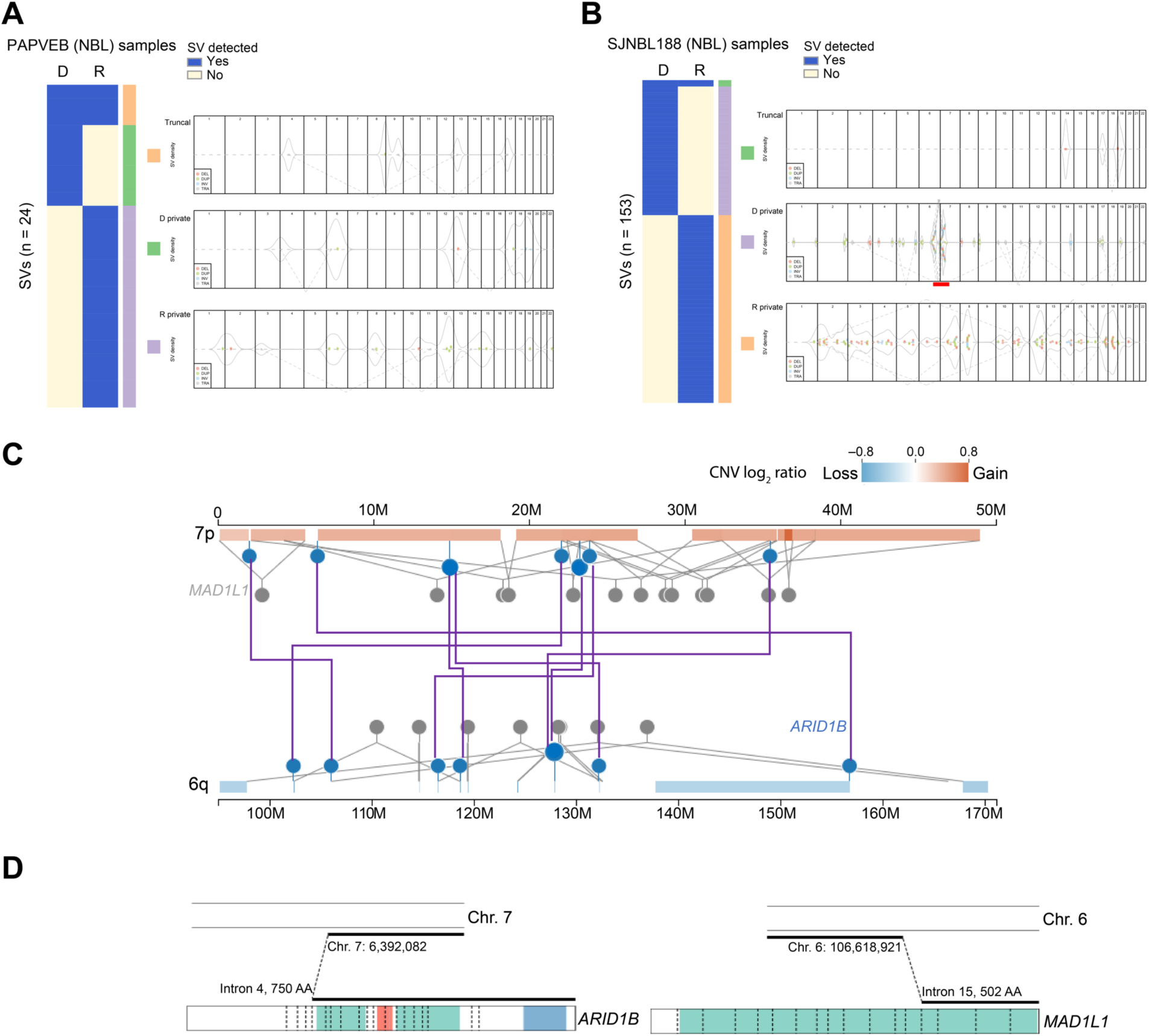
SV evolution in two multi-sample neuroblastoma patients. (**a**) SV grouping (left) and genome-wide density plot (right) shown in the same style as Figure 4 for two NBL samples obtained from one patient. D is from diagnosis while R is a relapse. (**b**) As in (a), except that a different patient is shown. D is from diagnosis while R is a relapse. The red bar at the 6q–7p region on the beeswarm plot for the SV group “D private” indicates a chromothripsis event. (**c**) Integrated view of SVs and CNVs at the 6q–7p region in (b) (region highlighted by a red bar) associated with a chromothripsis event targeting the *ARID1B* neuroblastoma driver gene. (**d**) Schematic detailing how this event disrupted both *ARID1B* and the common fragile site gene *MAD1L1*.

**Supplementary Figure 18.**
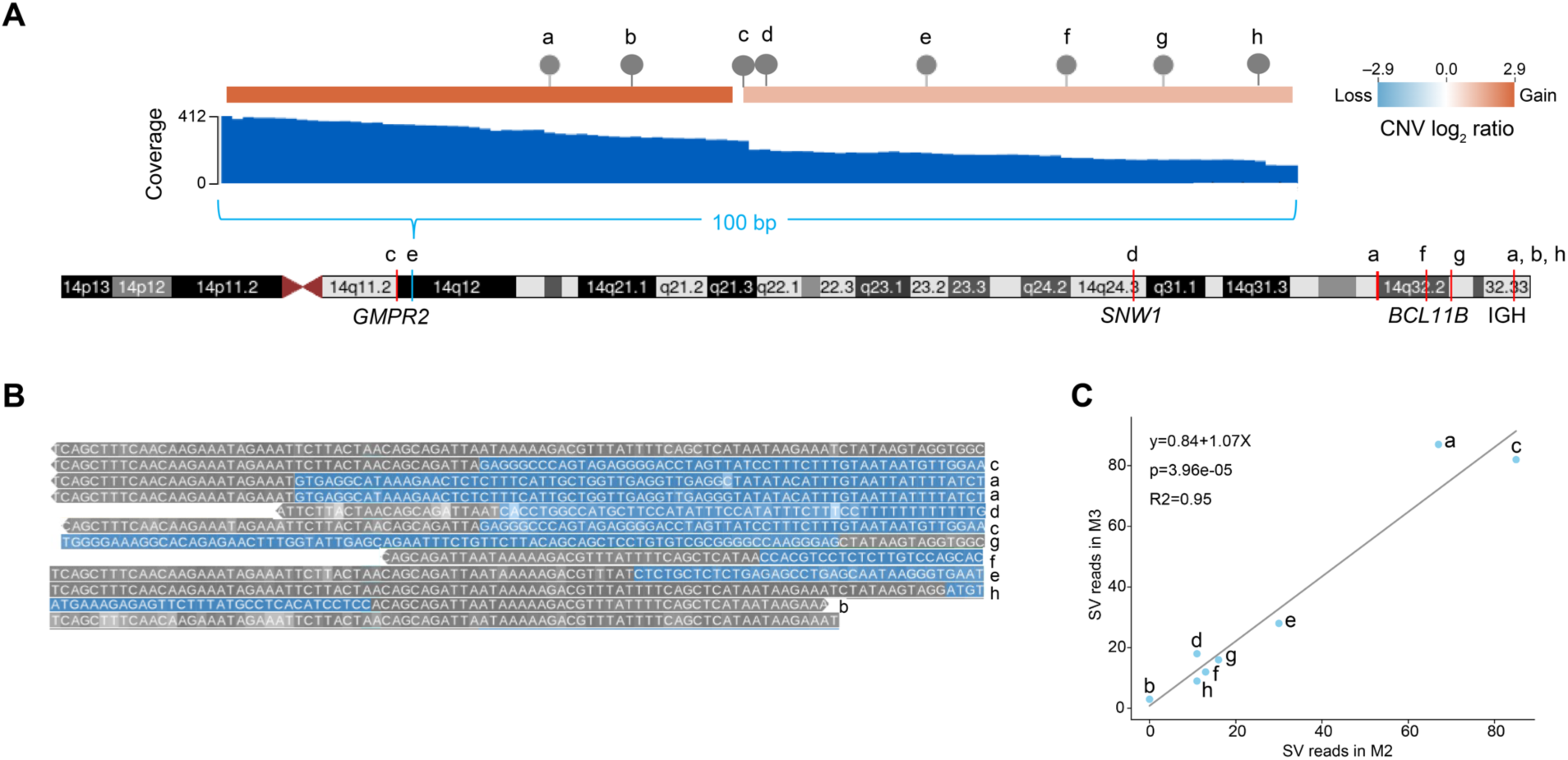
An amplicon with a high-density SV cluster defined as “shared” in multi-region analysis of osteosarcoma patient SJOS00101. This is a detailed view for the region shown in Figure 4b. (**a**) Eight SVs within 100 bp in an amplicon connected to seven distinct regions of chromosome 14. (**b**) Reference (top) and SV junction sequence (blue) at each of the eight breakpoints matching the labels in (a). (**c**) Concordant SV junction read counts between M2 and M3 samples. This suggests these may all be linked to a single, late event.

**Supplementary Figure 19.**
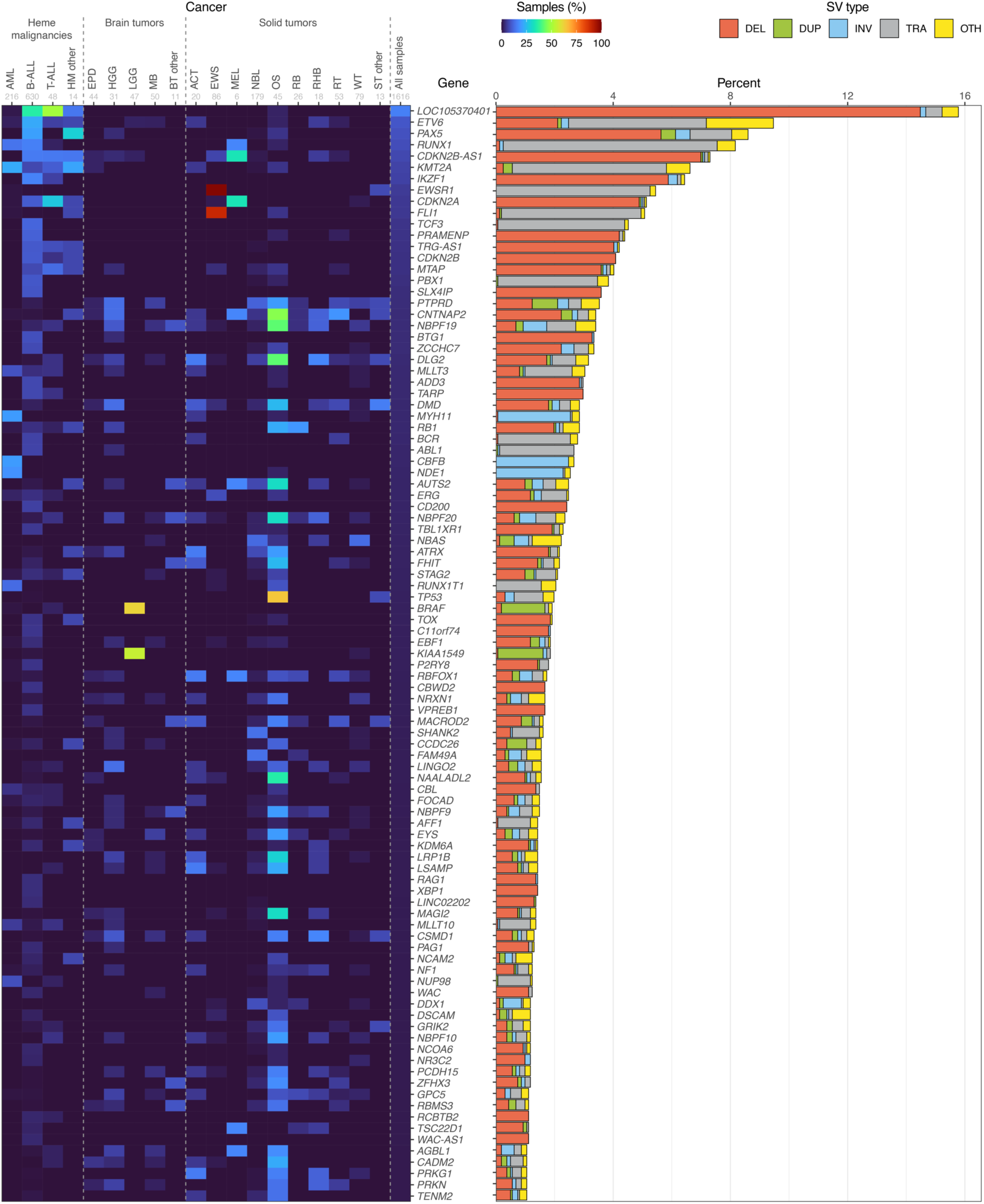
Frequency of intragenic SVs. Heatmap showing the percentage of samples in each pediatric cancer type (column) with somatic SVs impacting each gene (row). A gene was considered impacted in a sample if an SV breakpoint fell within any part of the gene including both exonic and intronic regions. Genes with SVs in at least 1% of pediatric cancer samples are shown and their order is sorted based on the prevalence of the entire cohort. Bar plot at right shows the percent of all pediatric cancer samples with SVs in each gene, separated by SV type. When multiple SV types within the same sample impacted the same gene, they are denoted with OTH.

**Supplementary Figure 20.**
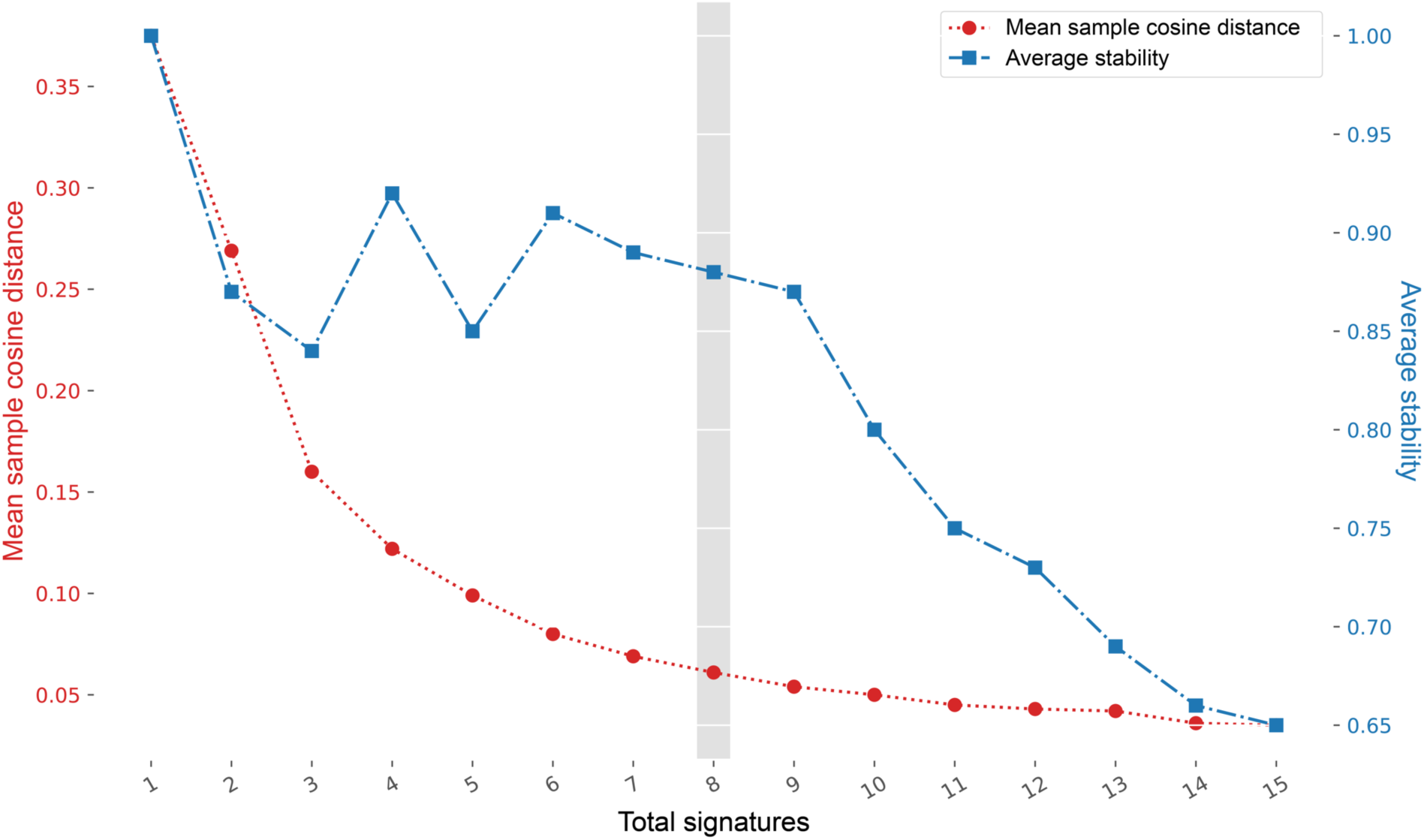
SigProfiler optimal selection plot for the pediatric cohort. Optimal number of signatures for the pediatric cohort as determined by `SigProfiler`. Left y-axis (red): Mean sample cosine distance; right y-axis (blue): average stability; x-axis: total number of signatures.

**Supplementary Figure 21.**
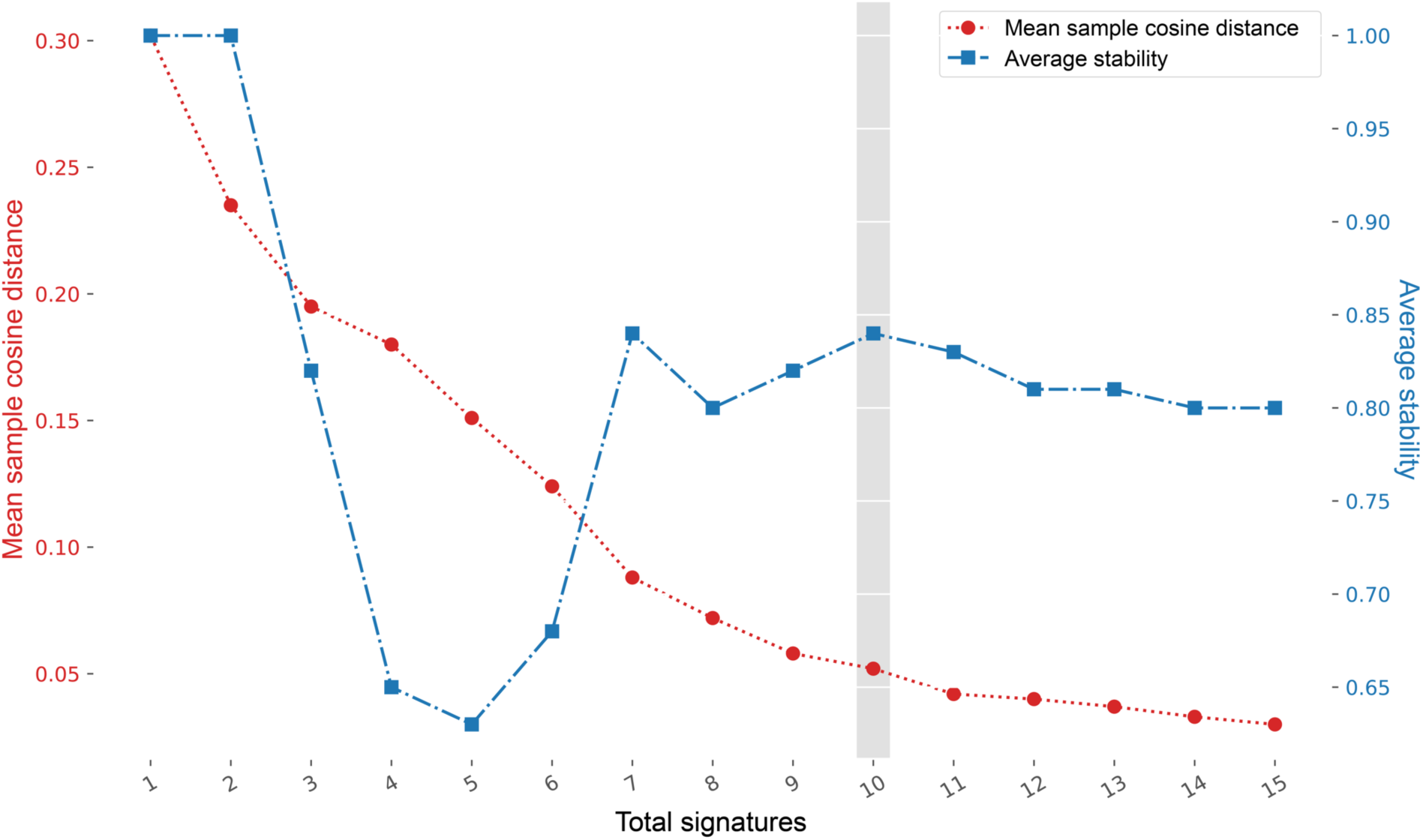
SigProfiler optimal selection plot for the adult cohort. Optimal number of signatures for the adult cohort as determined by `SigProfiler`. Left y-axis (red): Mean sample cosine distance; right y-axis (blue): average stability; x-axis: total number of signatures.

